# Metadomain and metaloop genome interactions in mammalian T cells

**DOI:** 10.1101/2025.09.19.677419

**Authors:** Gabriel Dolsten, Zhong-Min Wang, Xiao Huang, Susie Song, Michael J. Wilson, Xin Yang Bing, Wenfan Ke, Thomas R. Cafiero, Amy N. Nelson, Sebastian Fernando, Alexander Ploss, Paul Schedl, Michael S. Levine, Aaron D. Viny, Alexander Y. Rudensky, Yuri Pritykin

## Abstract

Recent studies have advanced our understanding of chromosomal organization and its principal role in gene regulation. However, most analyses have focused on short-range interactions (<2 Mb), limiting insight into broader regulatory architecture. In particular, the relationships between topologically associating domains (TADs), sub-TAD loops, long-range cross-TAD interactions, and higher-order chromosomal compartmentalization remain poorly understood. Here, we identify extensive multi-megabase and interchromosomal interactions (metaloops) in T lymphocytes, which organize into larger meta-TAD associations (metadomains). Metaloops bridge distal promoters and regulatory elements of key T cell-specific genes such as *Ctla4*, *Ikzf2*, *Il2ra*, *Ets1*, *Lef1*, *Runx1*, *Bach2*, *Foxo1* and others, and are both shared and cell type-specific across functionally distinct T cell lineages. Reanalysis of published data confirms the reproducibility of these interactions in both mouse and human T cells and their dependence on superenhancers. Genome-wide clustering of metadomains reveals three interchromosomal hubs with distinct epigenomic profiles, including a superenhancer-enriched hub associated with T cell-specific gene activation. By integrating a compendium of new and public T cell epigenomic data, we infer distinct architectural factors associated with short-range loops and long-range metaloops. Altogether, our study reveals new features of T cell-specific 3D genome organization across scales, and our computational framework is broadly applicable to analyses of chromatin architecture across different cell types and experimental systems.

**Highlights:**

- Ultra-long-range (>2 Mb) chromatin interactions linked to gene regulation in T cells
- A new algorithm identifies distal and interchromosomal meta-TADs and metaloops
- An interchromosomal T cell-specific active hub emerges from metadomain clustering
- Epigenomic compendium implicates TFs associated with long-range interactions

## Introduction

Advances in genomic, imaging, biochemical and computational methods have provided crucial insights into the spatial organization of chromatin (*1*–*6*). Chromosomal architecture in mammalian cells spans multiple scales. Chromatin contacts ranging from a few to several hundred kilobases (Kb) capture enhancer-promoter (E-P) interactions, CCCTC-binding factor (CTCF)- and cohesin-mediated loops, and interactions between tethering elements. Topologically associating domains (TADs), spanning 10 Kb to over a megabase (Mb), are regions where chromatin interactions are more frequent within the domain than with regions outside the domain, thereby constraining local genomic interactions and regulatory specificity. At the largest scale, A/B compartments segregate the genome into transcriptionally active (A compartment) and inactive (B compartment) regions, and chromosomes organize into territories (*7*). While these hierarchical features have been extensively described, an integrative model that links them across scales remains elusive, limiting our ability to dissect the relative contributions of local versus global chromatin organization to gene regulation. Recent findings further challenge the canonical hierarchy of genome architecture. In *Drosophila*, specific functional E-P metaloops spanning multiple megabases have been shown to cross TADs and compartment boundaries (*8*). In mammalian olfactory neurons, a specialized interchromosomal hub enables stochastic activation of a single olfactory receptor gene from among ∼20 “Greek island” loci dispersed across the genome (*9*, *10*). These examples suggest that E-P contacts can defy local topological constraints and assemble into large-scale spatially organized hubs, particularly in differentiated cells with stringent regulatory demands, such as neurons (*8*, *11*–*16*). However, data capturing such intermediate- and long-range interactions in physiologically relevant primary cells remain scarce, owing to cellular and tissue heterogeneity, limited cell numbers, and other confounders. Although several studies have characterized selected examples of functionally important long-range genomic interactions (*17*–*22*), there is a lack of data and tools for comprehensive genome-wide characterization of such features in functional primary cells within a spectrum of differentiation states and biological contexts.

Cells of the adaptive immune system offer unique opportunities for studying fundamental principles of gene regulation and chromosomal organization *in vivo* (*23*, *24*). Immune cells undergo activation, expansion, and diversification upon their differentiation into distinct, yet related and well-defined functional states, and undergo profound changes in gene expression during these processes. Additionally, they are amenable to highly efficient and stringent subset-specific isolation, aided by a rich armamentarium of genetic markers. In this study, we focused on two functionally opposing subsets of CD4^+^ T cells, “conventional” T (Tcon) cells that facilitate multifaceted pro-inflammatory immune responses, and regulatory T (Treg) cells, a specialized lineage with immunosuppressive function. The identity and function of Treg cells are defined by the highly restricted and stable expression of their lineage-specifying transcription factor (TF) *Foxp3*. Numerous studies have explored the regulatory genomics of Tcon and Treg cell differentiation and function (*25*–*28*). While recent studies have reported Treg cell-specific looping and chromatin remodeling and suggested a role of Foxp3 alongside other TFs in Treg chromatin organization (*29*–*33*), precise roles of these factors across different layers of chromatin organization remain incompletely understood, in part due to the limited resolution of existing Hi-C and HiChIP data and the lack of computational tools suited to characterizing intermediate- and large scale chromatin structures.

To address this, we developed a new computational pipeline for analysis of Hi-C data at large genomic scales and applied it to newly generated Hi-C data in Tcon and Treg cells. We first identified shared and cell type-specific TADs and local (< 2 Mb) genomic interactions at 5 Kb resolution and observed their association with gene expression. To investigate global features of chromatin architecture, we analyzed Hi-C data at 250 Kb resolution. We found that TADs surrounding certain cell type-specific genes had differential Hi-C interactions across entire chromosomes. To estimate the functional relevance of these interactions, we generalized the Activity-by-Contact (ABC) model of enhancer-promoter regulation and found that long-range (> 2 Mb) and interchromosomal interactions significantly contribute to cell type-specific gene expression regulation (*34*, *35*).

Thus, we set out to investigate features of chromosomal organization across different scales beyond conventional analysis of local loops, TADs, or A/B compartments. Our systematic analysis revealed thousands of multi-megabase-long and interchromosomal interactions between 250 Kb genomic regions, anchored by specific focal contacts between 5 Kb regulatory sites, similar to recently described metadomain and metaloop genomic interactions in Drosophila and the Greek islands in neurons (*8*, *9*). These interactions frequently connected promoters and enhancers of genes with a well-established functional importance in Treg and Tcon cells, including *Ikzf2*, *Ctla4*, *Cd28*, *Icos*, *Il2ra, Lef1*, *Ets1*, *Runx1*, *Bach2*, *Foxo1* and others. Reanalysis of published Hi-C and HiChIP data confirmed reproducibility of the metadomain interactions across studies in both mouse and human. Many metadomains linked T cell superenhancers (SEs), although not all SEs engaged in interactions with equal strength.

Genome-wide clustering of these interactions revealed multiple chromosome-wide and three distinct interchromosomal metadomain hubs. Each interchromosomal hub was enriched for specific active and repressive histone marks, suggesting they represent functionally and physically distinct subnuclear structures (*36*–*38*). While one active metadomain hub was associated with broad gene activation across many cell types, the other active hub was highly enriched for T cell-specific SEs and genes (*28*, *39*). Reanalysis of published Hi-C data following SE deletion demonstrated that this T cell-specific hub depended on SEs, suggesting that SEs may mediate long-range metadomain interactions (*18*). By building a compendium of public and newly generated Hi-C data across T cell subpopulations, we achieved 5 Kb resolution and identified E-P metaloop interactions spanning megabases. Integrative multi-omic analysis further implicated distinct regulatory factors in short-range looping versus long-range metalooping.

Together, these results suggest a heterogeneous landscape of functional chromatin interactions in primary T cells, with a high prevalence of focal enhancer-promoter loop and metaloop interactions and higher-order metadomain interactions occurring at all distance scales and between chromosomes. This work bridges our understanding of compartmentalization, transcriptional hubs, and local sub-TAD enhancer-promoter interactions. Our novel analytic approach is generally applicable to analysis of datasets across different cell types and experimental systems, and the associated software and results are shared via public repositories, to facilitate future research using these methods.

## Results

### Cell type-specific local and global chromatin organization in Tcon and Treg cells

To study chromatin architecture in T cells, we generated Hi-C data for Tcon and Treg cells isolated from the secondary lymphoid organs of healthy unchallenged female *Foxp3^GFP/LSL^* mice (**Figure 1A, Figure S1, S2**) (*40*, *41*). The Hi-C data were of high complexity, reproducibility and depth (**Figure S2, Table S1**), allowing us to identify 13,323 high-confidence local loops (< 2 Mb-long) at 5 Kb resolution (**Figure S2C-D, Table S2**). Of these, 10,636 loops were shared between Treg and Tcon cells, while 1,008 loops were significantly enriched in Treg cells (FDR < .05) and 1,679 in Tcon cells (**Figure S2E-G**). Many of the cell type-specific loops overlapped well-characterized genes differentially expressed in Treg cells such as *Ikzf2*, *Il2ra*, *Lrrc32* (*19*, *42*–*44*), as well as Tcon genes *Tgfbr3* and *Themis* (*45*–*48*) (**Figure 1B, Figure S2H-I, S3**). Thus, extensive differential chromatin looping occurs at many cell type-specific genes, underscoring the role of chromatin folding in regulating T cell identity and gene expression.

**Figure 1:**
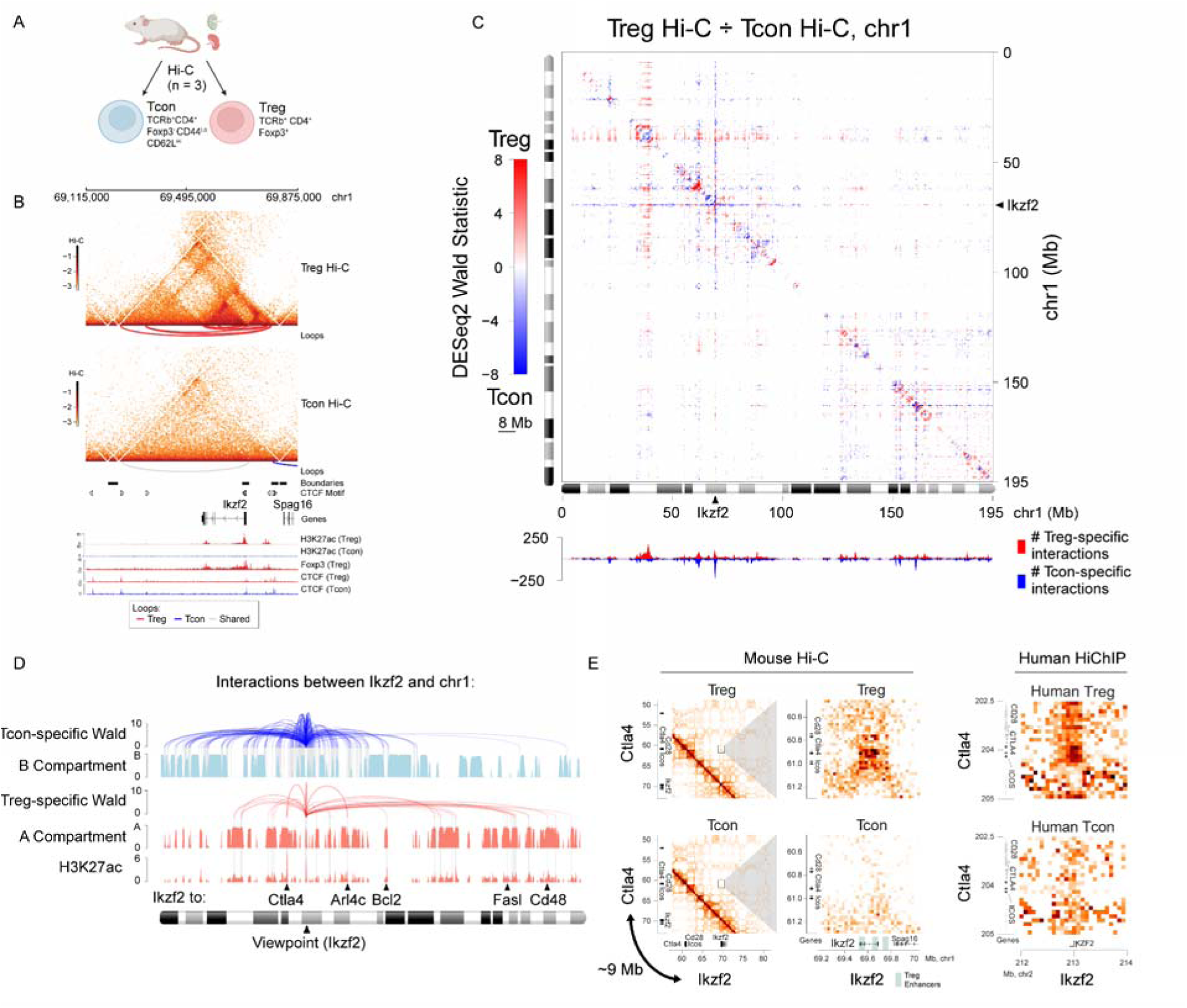
Cell type-specific local and global chromatin organization in Tcon and Treg cells. (A) Schematic of the experimental design. Hi-C (n = 3) was performed on Treg (TCRb+ CD4+ Foxp3(GFP)+) and Tcon (TCRb+ CD4+ Foxp3(GFP)-CD44^lo^ CD62L^hi^) cells isolated from the spleen and lymph nodes of healthy unchallenged female *Foxp3^GFP/LSL^* mice. (B) Balanced Hi-C chromatin contact frequency and epigenomic tracks at the *Ikzf2* locus. Red arcs: Treg-specific loops, blue arcs: Tcon-specific loops, gray arcs: shared loops. (C) Differential Hi-C analysis (DESeq2, see Methods) for 250 Kb genomic bins across chromosome 1. (Top) Wald statistic for Treg vs. Tcon comparison. Blue and red indicate significant differences (FDR < .05), otherwise white. (Bottom) Total number of significant Treg- (red) and Tcon-specific (blue) interactions. (D) Significant differential Hi-C interactions of *Ikzf2* along chromosome 1 (red and blue arcs) as in panel C, along with the A/B compartment scores in Treg cells (principal component loadings at 50 Kb resolution, see Methods), and Treg H3K27ac ChIP-seq signal aggregated over 50 Kb bins. (E) (Left) Balanced Hi-C contact map (25 Kb resolution) showing an interaction between *Ikzf2* and the locus encoding genes *Cd28*, *Ctla4* and *Icos* in our mouse Hi-C data. (Right) Balanced contact map (25 Kb resolution) of human CD4+ T cell H3K27ac HiChIP data (*17*) for interaction between *IKZF2* and the *CD28*/*CTLA4*/*ICOS* locus.

To assess how local genomic structural features relate to global chromosomal organization, we performed genome-wide A/B compartment analysis (*36*, *49*). We observed that anchors of Treg- and Tcon-specific loops had significantly different A/B compartment scores (**Figure S2J**), suggesting that differential local looping was associated with large-scale chromatin reorganization.

To further explore global chromosomal organization, we performed an unbiased genome-wide differential analysis of normalized Hi-C signal at 250 Kb resolution within each chromosome, approximating TAD-to-TAD interaction analysis (**Figure 1C, Figure S4, Table S3**). This analysis identified specific 250 Kb regions with significant reproducible differential Hi-C interactions across entire chromosomes (**Figure 1C, Figure S4C-E**). For example, the *Ikzf2* locus had 281 differential interactions across chromosome 1, including 157 interactions at distances > 20 Mb (**Figure 1G,H**). Genome-wide, 213 genomic loci, including *Lrrc32* and *Pde3b*, had more than 100 differential Hi-C interactions, and 46 loci, including *Ikzf2* and *Tgfbr3*, had more than 100 differential Hi-C interactions at distances > 20 Mb (**Figure S4E**). Remarkably, the Treg-specific gene *Ikzf2* formed many Treg-specific interactions with Treg-specific genes, including *Ctla4*, *Fasl* and *Arl4c*, which resided in the A compartment and were enriched for the active histone mark H3K27ac (**Figure 1D**). In addition, *Ikzf2* also engaged in many Tcon-specific interactions, particularly with regions less strongly associated with the A compartment and depleted of H3K27ac (**Figure 1D**). Notably, *Ikzf2* had a Treg-specific interaction, likely involving its promoter and enhancers, with a region 9 Mb away on chromosome 1 containing the genes *Cd28*, *Ctla4* and *Icos* which encode T cell co-stimulatory and co-inhibitory receptors (**Figure 1E)**. Reanalysis of published Hi-C and HiChIP data from both mouse and human confirmed reproducibility of this Treg-specific distal interaction (**Figure 1E, Figure S5A**). Overall, our analysis revealed a rich landscape of differential long-range interactions at 250 Kb resolution in Treg and Tcon cells, suggestive of TAD-to-TAD associations likely anchored by focal contacts between regulatory elements. These include cell type-specific interactions between functionally important T cell genes such as *Ikzf2*, *Ctla4* and others, highlighting the link between global 3D genome organization and cell type-specific gene regulation.

Therefore, we next sought to assess the functional relevance of distal chromatin interactions in Tcon and Treg cells. For this, we turned to the Activity-by-Contact (ABC) model of enhancer-promoter regulation (*34*, *35*). In this model, the regulatory potential of a putative enhancer is estimated as the product of its activity (e.g. H3K27ac ChIP-seq signal) and its contact frequency with a TSS (measured by Hi-C) (**Figure 2A**). Despite its simplicity, the ABC score is a strong predictor of enhancer activity for a gene (*34*, *35*). Therefore we reasoned that summing ABC scores over multiple regulatory elements of a gene could provide a measure of their collective regulatory contribution. To investigate the impact of distal chromatin interactions to gene expression, we developed a straightforward extension of the ABC model. For each gene, we calculated the cumulative ABC (cABC) score by summing ABC scores across all genomic positions located beyond 10 Kb from the TSS, within defined genomic windows around the TSS (e.g., 50 Kb, 1 Mb, entire chromosome, or genome-wide) (**Figure 2A**). We found that a substantial fraction of cABC signal was conferred by distal and interchromosomal elements (**Figure 2B**). Importantly, cABC scores were significantly correlated with gene expression across chromosomes, with the correlation strength increasing as the genomic distance cutoff expanded (**Figure 2C**). The strongest correlations were observed when cABC was calculated over entire chromosomes or the whole genome, indicating a significant contribution of distal and even interchromosomal interactions to gene expression. Similarly, differential cABC scores between Tcon and Treg cells also significantly correlated with differential gene expression (**Figure 2D**), suggesting that long-range interactions may help drive cell type-specific transcriptional programs. Only weak correlations were observed when using a computationally perturbed control (shifted H3K27ac signal), confirming biological relevance of the cABC scores (**Figure 2C,D**). Together, these results suggest that a substantial fraction of gene regulation could be attributed to long-range and interchromosomal interactions.

**Figure 2:**
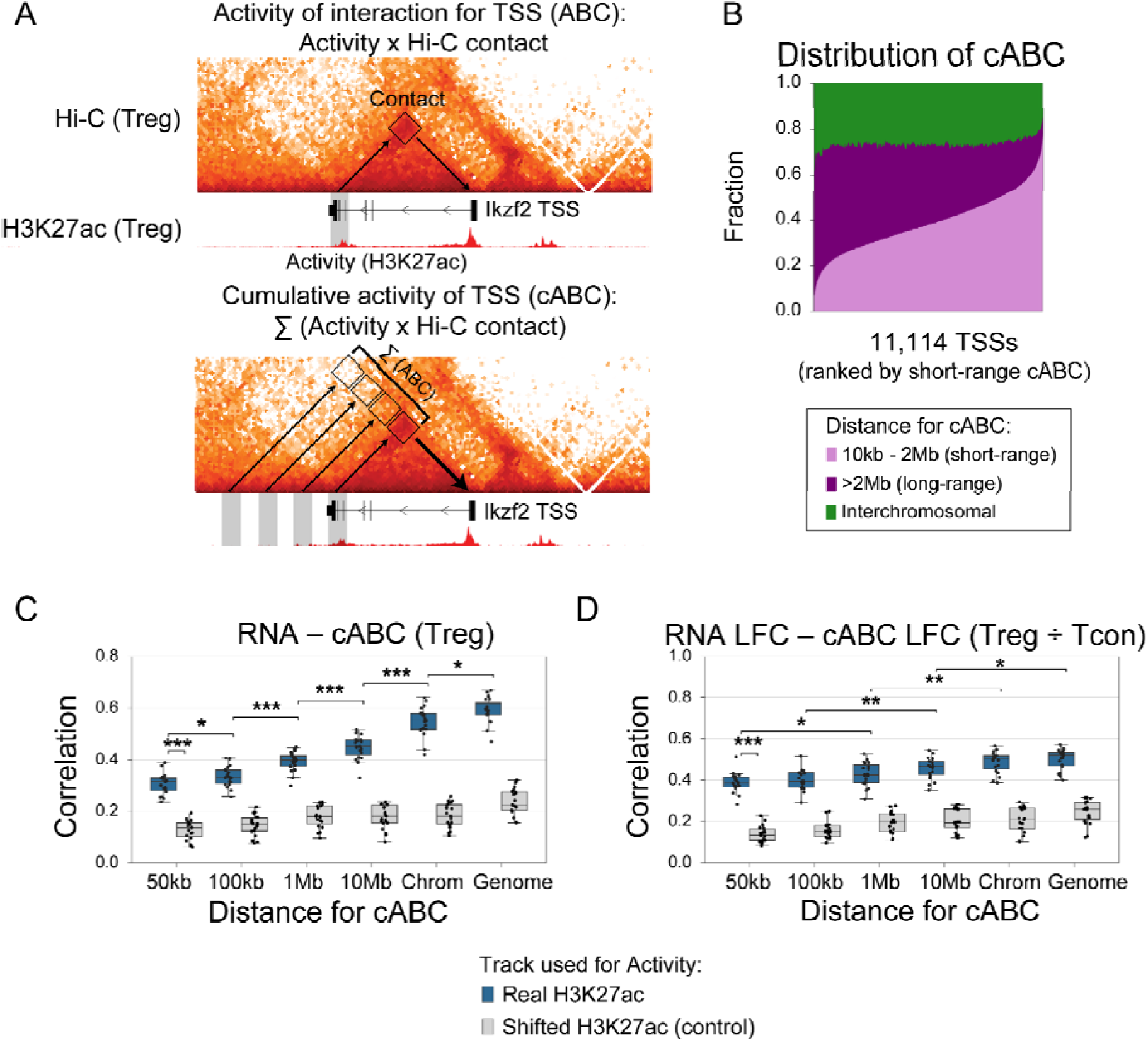
Cumulative activity-by-contact (ABC) analysis reveals the contribution of distal chromatin interactions to gene regulation in Tcon and Treg cells. (A) Schematic of the activity-by-contact (ABC) and cumulative activity-by-contact (cABC) score calculations. The ABC score of regulatory activity of a genomic site *S* for a transcription start site (TSS) is calculated as the product of the normalized Hi-C contact frequency between *S* and the TSS, and the activity of *S* represented by H3K27ac ChIP-seq signal. The cABC score for a TSS is calculated as the sum of ABC scores for this TSS over all genomic sites farther than 10 Kb from the TSS and within a defined distance window, e.g. within 50 Kb, 100 Kb, etc., or the entire chromosome, or the entire genome including interchromosomal contacts. (B) Stacked barplot of cABC scores conferred by interactions at varying distance cutoffs. (C) Pearson correlation between normalized gene expression (RNA-seq RPKM, reads per kilobase million) and cABC scores across genes on each chromosome, calculated at different distance cutoffs. As a control, the same calculation was performed for the H3K27ac ChIP-seq signal shifted 200 Kb to the right. Boxplots show the distribution of correlation values across chromosomes for 11,114 genes expressed in Tcon or Treg cells. **, p < .01; ***, p < .001. Boxplots: center line, median; box limits, upper and lower quartiles; whiskers, 1.5x interquartile range. (D) Pearson correlation between log2 fold change (LFC) in gene expression (Treg / Tcon) and LFC in cABC scores, calculated as in panel C. **, p < .01; ***, p < .001. Boxplots: center line, median; box limits, upper and lower quartiles; whiskers, 1.5x interquartile range.

### A new algorithm InterDomain identifies megabase-scale metadomain chromatin interactions

Having demonstrated the regulatory relevance of both local and global chromatin organization in Tcon and Treg cells, we next sought to systematically investigate specific long-range (> 2 Mb) interactions such as those between *Ikzf2* and the *Cd28*/*Ctla4*/*Icos* locus (**Figure 1E**). For this, we developed a new algorithm InterDomain by adapting and extending the HiCCUPS loop calling procedure (*38*) to enable the detection of long-range contacts (**Figure 3A**). InterDomain is designed for chromosome-wide Hi-C analysis at coarse resolution (e.g. 50 Kb genomic bins), and identifies regions with enriched contact frequency relative to their local background. After merging the signal across adjacent 50 Kb bins, the algorithm reports specific interactions at 250 Kb resolution. This approximates TAD-to-TAD metadomain interactions, or metadomains, analogous to how high-resolution local binned Hi-C loop calling approximates CTCF-mediated or enhancer-promoter loops. Using InterDomain, we identified 26,375 metadomain interactions between 250 Kb bins in the Treg and Tcon cell Hi-C data, representing .04% of all possible intrachromosomal 250 Kb bin pairs (**Figure 3A-D, Figures S5, S7, Table S4**). These metadomains captured interactions between many genes important for T cell function, such as between *Ikzf2* and the *Cd28*/*Ctla4*/*Icos* locus (9 Mb apart); *Ikzf2* and *Bcl2* (37 Mb); *Bcl2* and *Irf6* (86 Mb); *Izumo1r* and *Ets1* (18 Mb); and *Tox* and *Tgfbr1* (40 Mb) (**Figure 1E, 3B, Figure S5**). The metadomains spanned distances from 2-180 Mb (median 30 Mb, **Figure 3C**), and 80% of metadomains involved interactions within the A compartment (**Figure S7E**). Overall, our new algorithm detected thousands of reproducible megabase-scale intrachromosomal metadomain interactions in T cells, largely within active (A compartment) chromatin and involving numerous functionally important T cell genes.

**Figure 3:**
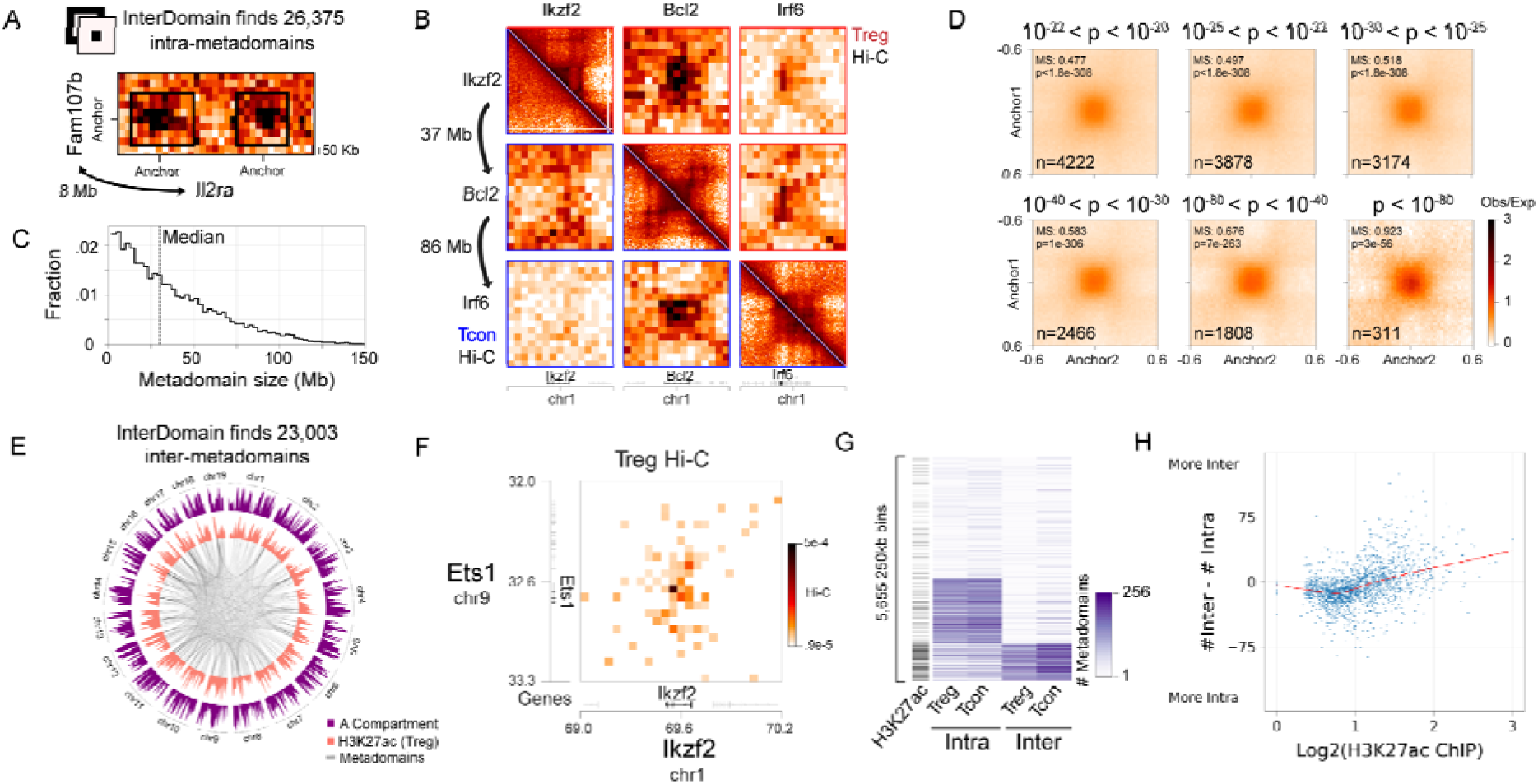
A new algorithm InterDomain identifies multi-megabase-scale and interchromosomal metadomain chromatin interactions in T cells. (A) InterDomain identifies 26,375 significant interactions in Tcon and Treg cells between 250 Kb genomic regions at distances greater than 2 Mb, called metadomains. The algorithm is applied to balanced Hi-C data at 50 Kb resolution. Shown are two examples of metadomains in Treg cells involving the genomic region near the Treg-specific gene *Il2ra*. (B) Balanced Hi-C contact maps (upper triangle, Treg; lower triangle, Tcon) showing a metadomain triplet spanning ∼120 Mb on chromosome 1, linking *Ikzf2*, *Bcl2* and *Irf6*. (C) Histogram of genomic distances between genomic regions linked by metadomains. The median distance is 30 Mb (dotted vertical line). (D) Pileup of log2(observed / expected) (log(O/E)) balanced Treg Hi-C (25 Kb resolution) for metadomains, stratified by InterDomain p-value (shown on top). (E) Circos plot displaying interchromosomal metadomain interactions identified by InterDomain (250 Kb bins; 50 Kb resolution Hi-C data input). The H3K27ac ChIP-seq signal and A/B compartment score are plotted along the periphery. Balanced Hi-C data (50 Kb resolution) showing an interchromosomal interaction between *Ets1* and *Ikzf2* in Treg cells. (G) Number of interchromosomal and intrachromosomal metadomains per 250 Kb genomic region. Clustering was performed using K-means clustering (K = 3). Average H3K27ac ChIP-seq signal is shown on the left (not used in clustering). (H) Scatterplot for 250 Kb bins with more than 10 intra- or interchromosomal metadomains. The x-axis shows Treg H3K27ac ChIP-seq signal; the y-axis shows the difference between the number of inter- and intrachromosomal metadomains per bin in Treg cells.

### InterDomain algorithm identifies interchromosomal metadomains

Given the substantial contribution of interchromosomal contacts to gene regulation (**Figure 2B-D**), we extended the InterDomain algorithm to interchromosomal Hi-C analysis and applied it to Tcon and Treg data. This analysis identified 23,003 reproducible interchromosomal metadomain interactions between 250 Kb genomic regions (**Figure 3E, Figure S6, S7, Table S4**). These interactions connected many important T cell genes, such as *Ikzf2* (chromosome 1) and *Ets1* (chr9), *Lef1* (chr3) and *Bach2* (chr4, a known cohesin-dependent gene (*50*)), *Cxcr5* (chr9) and *Ptprcap* (chr19), and *Jak1* (chr4) and *Itpkb* (chr1), and many of these interactions were likely centered at specific promoter-enhancer contacts, rather than reflecting broad compartmental colocalization (**Figure 3F, Figure S6**). Some loci were especially prolific, participating in 50 or more interchromosomal metadomain interactions (**Figure S7B**). While the number of inter- and intrachromosomal metadomains per bin was highly concordant between Treg and Tcon cells (**Figure 3G, Figure S7C**), distinct sets of bins were preferentially involved in intra- or interchromosomal interactions. These categories were associated with intermediate and high levels of H3K27ac signal, respectively (**Figure 3G, Figure S7D**). For instance, on chromosome 1, both *Ly96* and *Ikzf2* were involved in numerous intrachromosomal metadomains, but only *Ikzf2* showed extensive interchromosomal interactions (**Figure S6C-D**). Thus, in addition to intrachromosomal metadomains, InterDomain enabled systematic identification of interchromosomal metadomain interactions at 250 Kb resolution, many of which connected genes critical for T cell activation and function.

In sum, our analysis revealed that metadomains, representing a previously underappreciated feature of chromosomal organization in T cells beyond local TADs and global A/B compartments, are widespread throughout the genome with potential for gene regulatory function.

### Interchromosomal metadomains segregate into one repressive and two distinct active hubs

To investigate higher order genome organization and functionally characterize different classes of metadomains, we clustered all intra- and interchromosomal metadomains identified at 250 Kb resolution. While most clusters were intrachromosomal, eight of the clusters were predominantly interchromosomal, involving from 14 to 20 different chromosomes (**Figure S8A, Table S4**). Some interchromosomal clusters were enriched for specific histone marks, including H3K27ac, H3K4me1, H3K4me3 and H3K27me3, while intrachromosomal clusters showed no such enrichment (**Figure S8B**). Thus we identified multiple intra- and interchromosomal metadomain clusters with distinct active and repressive epigenetic signatures, with metadomain analysis providing a framework for defining higher order features of chromosomal organization.

Interchromosomal metadomain clusters were merged into three interchromosomal hubs. Based on chromatin and transcriptional features, we annotated these hubs as Active Constitutive, Active Dynamic, and Repressive (**Figure 4, Figure S8, S9**). Both active hubs were highly enriched for superenhancers (SEs), together encompassing ∼50% of all T cell SEs (**Figure S8F**). The Active Constitutive hub was specifically enriched for the active promoter mark H3K4me3 and for housekeeping genes broadly expressed across many cell types, including T cells (**Figure 4D, E, G**). In addition, it was also associated with nuclear speckles, as evidenced by significant proximity to speckles in mouse embryonic stem cells (mESCs), and had a high gene density, and these genes were often short and exon-dense, consistent with speckle-associated genomic regions (**Figure S9E**) (*52*, *53*). Analysis of mESC Hi-C data (*54*) confirmed the presence of this hub in mESCs (**Figure S8C, S9K**), suggesting that it represents a constitutive feature of genome organization. Notably, although broadly constitutive, the hub also included several T cell-specific genes such as *Id3*, *Lck*, *Lag3*, *Cxcr5* and *Il10*, indicating potential cell type-specific plasticity.

**Figure 4:**
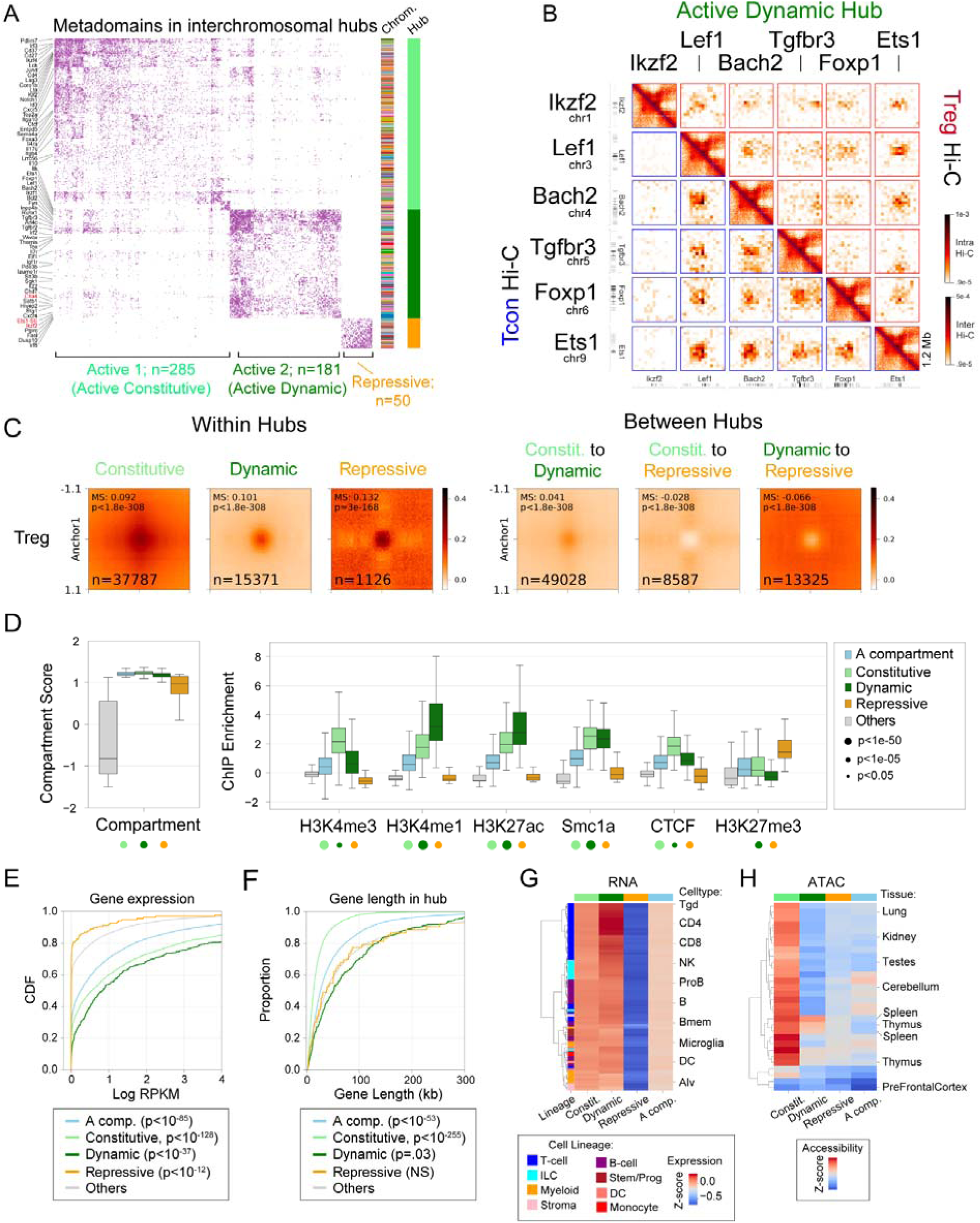
Interchromosomal metadomain interactions segregate into three hubs with distinct epigenomic profiles. (A) Visualization of inter-chromosomal hubs formed by 250 Kb genomic bins involved in metadomain interactions identified by InterDomain. Hubs are categorized as Active 1 (Constitutive), Active 2 (Dynamic), and Repressive. Hub assignments and chromosome identities (indicated by unique colors) are shown on the right. (B) Balanced Hi-C contact maps (50 Kb resolution; upper triangle, Treg; lower triangle, Tcon) for selected genes within the Active 2 (Dynamic) interchromosomal hub. (C) Aggregate interchromosomal Hi-C signal (25 Kb resolution) in Treg cells within (left) and between (right) the metadomain hubs. (D) A/B compartment score (left) and normalized ChIP-seq signal (right) aggregated over 250 Kb bins within each metadomain hub. The A compartment bins serve as a control (statistical comparisons against the A compartment bins, Mann-Whitney U test). Boxplots: center line, median; box limits, upper and lower quartiles; whiskers, 1.5x interquartile range. (E) Cumulative distribution function (CDF) plot for the normalized gene expression values (Treg RNA-seq RPKM) for genes in interchromosomal metadomain hubs, A compartment and other genomic regions (statistical comparisons against “Others”, Mann-Whitney U test). (F) CDF plot of the gene lengths for genes in interchromosomal metadomain hubs, A compartment and other genomic regions (statistical comparisons against “Others”, Mann-Whitney U test). (G) Aggregated normalized gene expression across immune cell types (ImmGen Consortium data) for genes in interchromosomal metadomain hubs. RNA-seq log(RPKM) values were z-scored across genes within each cell type; the z-scores for hub-associated (or A compartment) genes were then averaged to calculate hub-level activity. (Selected cell types are labeled: Tgd, gamma-delta T cells; CD4, CD4 T cells; CD8, CD8 T cells; NK, natural killer cells; ProB, progenitor B cells; B, B cells; Bmem, memory B cells; DC, dendritic cells; Alv, alveolar macrophages; ILC, innate lymphoid cells; Stem/Prog, stem and progenitor cells.) (H) Aggregated chromatin accessibility (ATAC-seq) across tissues and cell types (data from (*51*)) for genes in interchromosomal metadomain hubs. ATAC-seq values were z-scored per cell type, and hub- or A-compartment-associated gene values were averaged to calculate gene activity for each hub. Selected tissues and cell types are labeled.

In contrast, the Active Dynamic hub was enriched for enhancer-associated histone marks such as H3K4me1 and H3K27ac, suggestive of dynamic regulatory activity at these sites (**Figure 4B,D**). While the Constitutive hub genes were broadly expressed across cell types, the Dynamic hub genes showed predominant expression in lymphocytes and T cells, and included many well-established T cell-specific genes such as *Ikzf2*, *Ikzf1*, *Lef1*, *Ets1*, *Tox*, *Runx1*, *Irf2*, *Irf6*, *Bcl2*, *Foxp1* and *Ctla4* (**Figure 4E-H**). Notably, this hub did not form even weak contacts in mESCs, confirming its cell type-specificity (**Figure S8C, S9K**). While ATAC-seq peaks in the Constitutive hub were broadly accessible across tissues and organs, those in the Dynamic hub showed spleen- and thymus-specific accessibility and were enriched with TF motifs with a prominent activity in T cells, including Ets, Stat, Irf and Tcf (**Figure 4H, Figure S9J**), suggesting establishment during thymic T cell differentiation. Among the three hubs, only the Dynamic hub was enriched for ATAC-seq peaks differentially accessible between Treg and Tcon cells, underscoring its role in cell type-specific gene regulation within the T cell lineage (**Figure S9H**). Unlike genes in the Constitutive hub, genes within the Dynamic hub were longer on average and had lower exon density, features that are inconsistent with those typically associated with nuclear speckles (**Figure 4F, Figure S9D**). Thus, we identified two distinct active interchromosomal hubs, each marked by unique epigenomic signatures and spatially segregated into specialized nuclear domains, potentially reflecting unique regulatory and organizational requirements. The Active Constitutive hub is associated with constitutive gene expression across cell types and organs, and the Active Dynamic hub with T cell-specific gene regulation.

The Repressive hub was enriched for the polycomb-associated repressive histone mark H3K27me3. While the Active hubs showed some degree of inter-hub contact, the Repressive hub was markedly depleted for interactions with either Active hub, suggesting spatial segregation into a distinct nuclear compartment (**Figure 4C, Figure S8C**). Genes within the Repressive hub were consistently lowly expressed across multiple cell types (**Figure 4E,G**), indicative of constitutive transcriptional repression. Interestingly, the Repressive hub was also associated with nuclear speckles in mESCs, suggesting that it represents a repressive subset of nuclear speckles physically segregated from active speckle-associated regions (**Figure S9E**) (*52*). Although all three interchromosomal hubs resided in the A compartment, the Repressive hub had significantly lower A compartment scores than either Active hub, suggesting a partial or intermediate A compartment state (**Figure 4D, Figure S9B**). Moreover, the majority of A-compartment bins, many with strong A scores, were not part of any of the interchromosomal hubs (**Figure S8G**). Clustering of the normalized Hi-C signal at the same 250 Kb resolution, but without metadomains, recovered these hubs, demonstrating the robustness of our classification (**Figure S8E**). Previously defined subcompartments did not show a specific association with our interchromosomal metadomain hubs, for example all three hubs overlapped subcompartment A1 to a similar degree as other bins with high A compartment score (**Figure S9I**) (*55*). Thus the interchromosomal metadomain hubs represent a robust and novel feature of global chromosomal organization in T cells, distinct from classical A/B compartments or subcompartments.

### Cell type specificity of metadomain hubs and association with superenhancers

Having established the existence of distinct interchromosomal hubs in Tcon and Treg cells, next we sought to further investigate their functional properties, dynamics and cell type specificity.

Given that the Active Dynamic hub appeared T cell specific, we asked when during thymic development this hub first emerged. To address this, we reanalyzed published staged Hi-C data across thymic differentiation (*29*) and quantified aggregated contact strength within each of the three hubs, as well as the A compartment (**Figure S8C**). All three hubs showed a general increase in contact strength over the course of differentiation (**Figure 5A**). Notably, the Dynamic hub was largely absent at the double-negative (DN) stage, showing contact levels comparable to the baseline levels of other bins with similar A compartment strength (**Figure 5A,B**). In contrast, significant enrichment relative to these baseline levels was observed beginning at the double-positive (DP) stage and persisted through later stages (**Figure 5A,B**). These results suggested that the T cell-specific Active Dynamic hub is established at the DP stage and continues to strengthen as T cells transition from a quiescent to a more transcriptionally active state during thymic differentiation (*56*).

**Figure 5:**
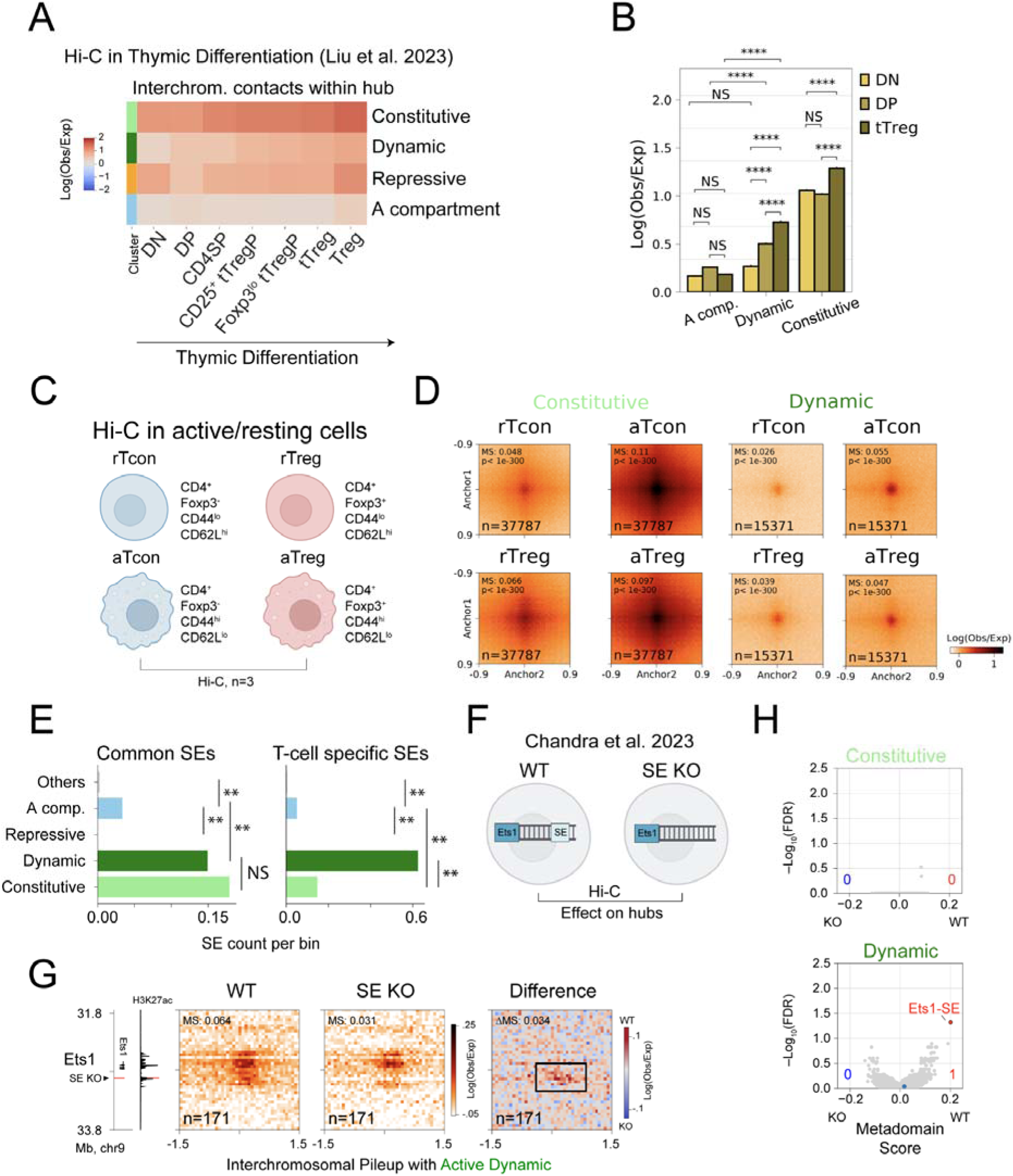
Cell type specificity of metadomain hubs and association with superenhancers. (A) Average interchromosomal log2(observed / expected) Hi-C contact frequencies for metadomain hubs across thymic differentiation. (DN, double-negative CD4^-^ CD8^-^ T cells; DP, double-positive CD4^+^ CD8^+^ T cells; CD4SP, single-positive CD4^+^CD25^-^ CD8^-^ T cells; CD25^+^ tTregP, CD4^+^ CD8^-^ CD25^+^ Foxp3^-^ T cells; Foxp3^lo^ tTregP, CD4^+^ CD8^-^ CD25^+^ Foxp3^lo^ T cells; tTreg, CD4^+^ CD8^-^ CD25^+^ Foxp3^+^ cells. Data from Liu et al. 2023 (*29*).) (B) Quantification of signal in panel A for DN, DP and tTreg cells for both Active hubs and the A compartment. (****, p < 1e-5, Mann-Whitney U test, with ΔLog(Obs/Exp) > .2) (C) Schematic of Hi-C experiments (n = 3) performed in activated (CD44^hi^ CD62L^lo^) and resting (CD44^lo^ CD62L^hi^) Tcon and Treg cells. (D) Aggregated interchromosomal Hi-C signal (25 Kb resolution) in activated and resting Tcon and Treg cells for the Active Constitutive and Active Dynamic metadomain hubs. (E) Frequency of common superenhancers (SEs; shared between mouse embryonic stem cells and T cells) or T cell-specific SEs overlapping 250 Kb genomic bins in each interchromosomal hub. (**, p < 1e-3, Fisher’s Exact Test) (F) Schematic of Hi-C experimental design from Chandra et al. 2023 (*18*). Wildtype (WT) and partial *Ets1*-SE knockout (Ets1-SE KO) CD4 Th1 cells were profiled by Hi-C. We reanalyzed this dataset to assess the impact of *Ets1*-SE deletion on metadomains in T cells. (G) Aggregate interchromosomal Hi-C signal (25 Kb resolution) between the *Ets1* locus and the Active Dynamic hub in WT (left) or *Ets1*-SE-KO (right) Th1 cells. The *Ets1*-SE overlaps a single 25 Kb bin, highlighted in red. (H) Scatterplot showing the metadomain score for interactions between each 250 Kb genomic bin and the Active Constitutive (top) or Active Dynamic (bottom) hub. A modified flanking region was used to calculate differential signal, accounting for global signal reduction following Ets1-SE KO (see Methods). The single genomic region including Ets1-SE with significantly altered interaction score (FDR < 0.05) is shown in red.

While metadomain hubs were established at least as early as during thymic differentiation and were broadly shared between Tcon and Treg cells (**Figure 4C, Figure S7C-F**), they were also differentially associated with Tcon-specific and Treg-specific SEs (**Figure S10F**), suggesting potential cell type specificity. Certain loci, such as *Ikzf2* or *Tgfbr3*, also showed focal cell type-specific interactions (**Figure 4B**). Therefore, we wanted to explore the level of plasticity and cell type specificity of metadomains in Treg and Tcon cells and their association with gene expression.

Focusing only on the interchromosomal Hi-C signal, we identified genomic regions with differential aggregate contact with each metadomain hub in Treg compared to Tcon cells (**Figure S8C, S10A**). Genomic regions with cell type-specific interactions with the Active Constitutive and Active Dynamic hubs were associated with increased gene expression in the corresponding cell type, whereas genes with stronger contact with the Repressive hub showed decreased gene expression (**Figure S10B**). Some loci gained Treg- or Tcon-specific contacts in a hub-specific manner. For example, *Ikzf2* and *Ctla4* both had differential contact with the Dynamic hub, but not the Constitutive hub (**Figure S10C,D**). Among these, *Ikzf2* demonstrated the strongest Treg-specific involvement in the Dynamic hub (**Figure S10C**). These findings highlight that, despite the overall similarity of metadomains and metadomain hubs between Tcon and Treg cells, substantial cell type-specific differences in hub connectivity exist, with *Ikzf2* serving as a prominent example of a Treg-specific locus engaging in the Active Dynamic hub.

T cells require coordinated upregulation of gene expression during activation in response to immunological challenges, and interchromosomal hubs may support this regulation. To assess this, we analyzed gene co-expression within the three hubs using published Treg scRNA-seq data (*57*). Genes within the same active hub were significantly more co-expressed than genes outside the hub, indicating coordinated expression of genes that are distal in genomic coordinates but share interchromosomal environments (**Figure S10E**) (*58*, *59*). Notably, while the Active Constitutive hub showed modest but significant co-expression, the T cell-specific Active Dynamic hub exhibited substantially stronger coordination. These findings suggest that interchromosomal hub formation may facilitate regulation of T cell gene programs.

To further examine the relationship between interchromosomal hubs and T cell activation, we generated Hi-C data separately for activated (CD44^hi^ CD62L^lo^) and resting (CD44^lo^ CD62L^hi^) Treg (aTreg, rTreg) and Tcon (aTcon, rTcon) cells (**Figure 5C**) (*40*, *41*, *60*). Within both Tcon and Treg cells, metadomain hubs were more pronounced in activated rather than in resting cells, suggesting that metadomain hubs were associated with T cell activation and thus higher transcriptional activity (**Figure 5D, Figure S10G**). Interestingly, even in the resting state, Treg cells exhibited stronger hub interactions than Tcon cells, suggesting a continuum of cell type-specific and immune activation-dependent chromatin organization (**Figure 5D, S10G**).

Moreover, Treg-specific involvement of *Ikzf2* in the active metadomain hubs was observed both for resting and activated Treg cells, whereas in Tcon cells, this interaction between *Ikzf2* and the Active Dynamic hub was only observed in the activated state (**Figure S10H,I**). Together, these results suggest that metadomains are both cell type-specific and associated with T cell activation.

To better understand the regulatory mechanisms of cell type-specificity and plasticity of metadomains and metadomain hubs, we examined the potential role of SEs in mediating their formation, given the established involvement of SEs in transcriptional hubs (*61*–*63*). We already demonstrated that both active metadomain hubs were enriched for SEs, and that metadomains were differentially associated with Tcon-specific and Treg-specific SEs (**Figure S8F, S10F**). To further explore this, we classified all T cell SEs based on whether they were shared with mESCs or T cell-specific. While both the Active Constitutive and Active Dynamic hubs were enriched for SEs, the Dynamic hub exhibited significantly higher enrichment for T cell-specific SEs (**Figure 5E**). We therefore hypothesized that different classes of SEs may selectively mediate interactions with the Dynamic and Constitutive hubs. To test this, we reanalyzed recently published Hi-C data from CD4 Th1 cells in which a SE near the T cell-specific gene *Ets1* was partially deleted (**Figure 5F**) (*18*). Both *Ets1* and *Ets1*-SE reside in the Active Dynamic metadomain hub (**Figure 4A**). Aggregate contact analysis revealed that the *Ets1*-SE deletion selectively impaired the interactions between the *Ets1*-SE locus and the Active Dynamic hub, while the contacts with the Active Constitutive hub remained unchanged (**Figure 5G,H**). Notably, the disruption extended beyond the precise site of *Ets1*-SE deletion, indicating that the *Ets1*-SE was required to maintain broader regional interactions within the hub, although the *Ets1* locus itself was not significantly affected. These findings indicate that the T cell-specific Active Dynamic hub depends on the *Ets1*-SE, suggesting that SEs may contribute to the formation of interchromosomal metadomain hubs.

Overall, our analysis reveals that metadomain hubs are dynamic, cell type- and immune activation-specific features of genome organization in T cells. We propose that these structures, established in part during thymic differentiation, support two modes of gene regulation via distinct higher-order nuclear structures: constitutive expression of broadly active genes in the Active Constitutive hub, and coordinated activation of T cell-specific programs in the Active Dynamic hub. In particular, we identified a previously undescribed interchromosomal hub that is associated with T cell-specific gene activation and dependent on T cell-specific SEs. These findings, together with the observed enrichment of cell type-specific SEs and plasticity in hub connectivity, suggest the importance of regulatory elements in organizing interchromosomal architecture.

### Comprehensive T cell Hi-C data compendium reveals enhancer-promoter interactions at 5 Kb resolution

We have shown that long-range metadomain interactions, likely anchored at specific enhancer-promoter (E-P) contacts, are closely associated with SEs, and our cABC analysis suggested that long-range interactions contribute to gene activity (**Figure 1E, 2, 3F, 5E-H, Figure S5B**). However, the inability to enhance the resolution of metadomain analysis beyond 250 Kb for a single dataset impedes detailed understanding of regulatory features of metadomains. To overcome this limitation, we reasoned that resolution could be enhanced by aggregating related datasets. Given the high degree of similarity in chromatin architecture between Tcon and Treg cells and their thymic precursors (**Figure 4B, 5A, Figure S2B,E, S7C-F**), we expected that many T cell subpopulations would share features of chromosomal organization across scales. Therefore, we aggregated 51 published T-cell Hi-C datasets, as well as newly generated high quality CD4 and CD8 T cell Hi-C data, to create a high-resolution T cell Hi-C compendium of unprecedented depth with 19 billion Hi-C contacts (**Figure S11, Table S4**). Visualization of these data confirmed punctate megabase-spanning interactions at 5 Kb resolution (**Figure 6A**). To identify specific genomic sites driving metadomain interactions, we applied InterDomain to this compendium at 5 Kb resolution, searching within the long-range metadomains detected in Tcon/Treg cells at 50 Kb resolution. We identified 26,037 focal interactions within metadomains, similar to recently described metaloops (*8*) (**Figure 6A,B, Figure S11, S12**). These metaloop interactions were significantly more frequent in metadomains than in the flanking regions (**Figure S11D**). By definition, these metaloops connected 5 Kb genomic regions separated by megabases, and included striking examples of E-P and promoter-promoter (P-P) loops between T cell-specific genes: for example, an 8Mb-long E-P contact between *Il2ra* and an intronic enhancer of *Fam107b*, and metaloops linking *Ikzf2* and *Idh1*, *Cd28* and *Stk16b,* and *Izumo1r* and *Birc3* (**Figure 6B, Figure S12**). Most metaloop anchors overlapped short-range loop anchors identified in Tcon and Treg cells, indicating that many long-range metaloops involved the same regulatory loci that mediate local E-P contacts (**Figure S11E**). These anchors frequently overlapped TSSs of highly expressed genes and were enriched for H3K27ac, consistent with high regulatory activity (**Figure S11G,H**). Metadomains within the Active hubs harbored significantly more metaloops than those in the Repressive hub, suggesting that Polycomb-associated repressive domains may primarily reflect compartmentalization rather than direct looping (**Figure 6C**). To further connect this high-resolution T cell Hi-C map with Treg- and Tcon-specific regulatory activity, we leveraged published Treg single-cell ATAC-seq data, which offers finer resolution than Hi-C, to examine co-accessibility of genomic regions shown to approximate chromosomal interactions (63–65). Metaloop anchors within metadomains showed significantly greater co-accessibility than random pairs of matched anchors, supporting the presence of these long-range interactions in Treg cells (**Figure 6D**). Thus, our T cell Hi-C compendium analysis revealed that metadomains harbor punctate 5 Kb metaloops, many of them likely E-P contacts, analogous to short-range regulatory loops.

**Figure 6:**
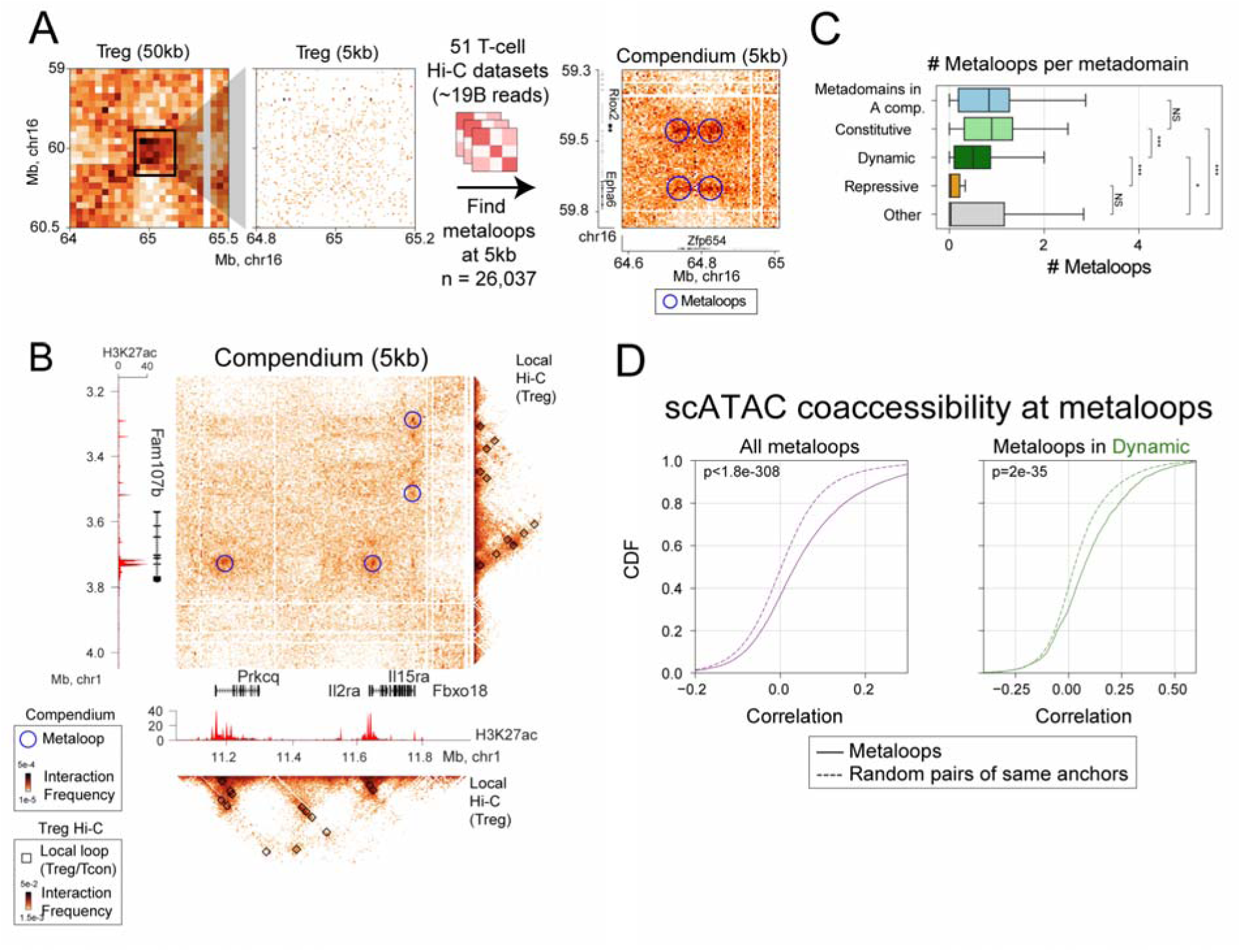
A comprehensive T cell Hi-C compendium reveals high-resolution metaloop interactions underlying metadomains. (A) Published and newly generated Hi-C data from T cells were combined into a compendium (51 samples, 19 billion processed contacts). This enabled the identification of distal focal interactions at 5 Kb resolution within intrachromosomal metadomains, called metaloops. (B) (Center) Balanced T cell Hi-C compendium data at 5 Kb resolution for the interactions between the *Il2ra* and *Fam107b* genomic regions that are separated by 8 Mb. Metaloops are indicated by circles. (Bottom and right) Balanced Treg Hi-C data at 5 Kb resolution for the *Il2ra* (bottom) and *Fam107b* (right) genomic regions. Local loops at 5 Kb resolution are indicated with squares. (C) Boxplot of the distribution of the number of metaloops per metadomain for different categories of genomic regions. Boxplots: center line, median; box limits, upper and lower quartiles; whiskers, 1.5x interquartile range. Mann-Whitney U test. (D) Single-cell ATAC-seq co-accessibility between metaloop anchors, compared to random pairs of the same anchors. The analysis was performed for metaloops genome-wide (left) and for metaloops in the Active Dynamic hub metadomains (right). Mann-Whitney U test.

### Integrative ChIP-seq, CUT&RUN and ATAC-seq data analysis implicates metaloop-associated regulatory elements and transcription factors in Tcon and Treg cells

After refining metadomains to putative 5 Kb focal metaloops, we wanted to search for regulatory drivers of metalooping. Therefore, we collected an epigenomic compendium combining 201 ChIP-seq, CUT&RUN, and ATAC-seq datasets spanning eight different Tcon and Treg cell studies (**Figure 7A,B, Table S5**). This resource enabled a systematic comparison of differential TF binding and histone modifications between anchors of long-range metaloops and short-range loops (**Figure 7C,D**). We observed that Stat5, Tet2, H3K4me3 and H3K27ac ChIP-seq signals were the most strongly enriched in long-range metaloop anchors, while H3K27me3, CTCF and cohesin (Smc1) were most strongly enriched in anchors of short-range loops (**Figure 7D,E**). Thus, known chromatin looping factors CTCF and cohesin were unlikely to be associated with long-range chromatin interactions, consistent with an estimated processivity of cohesin of 1-2 Mb (*64*, *65*). Notably, Foxp3 binding was not associated with Treg-specific looping, and was only weakly associated with loop anchors compared to other genomic regions, consistent with the finding that Foxp3 binds primarily to pre-existing T cell enhancers (**Figure S13B-D**) (*26*). As expected, H3K27ac, a mark of active enhancers and promoters, was enriched at long-range contact anchors, in line with prior results showing an association between long-range interactions and active regulatory elements, including SEs (**Figure 3A**). We therefore hypothesized that TFs associated with H3K27ac marks could mediate metadomains and metaloops. Among all factors examined, Treg Stat5 binding was most strongly correlated with Treg-specific H3K27ac signal (**Figure 7F,G**), Treg-specific looping (**Figure S13C-D**), and Stat5 motifs were among the most enriched motifs in Treg-specific H3K27ac peaks (**Figure 7H, Figure S13A**). Moreover, Stat5 binding was among the top correlates of Treg-specific contact with the Active interchromosomal hubs (**Figure 7I**). Thus, our analysis suggested that Stat5 may be associated with long-range genomic interactions in Treg cells. Overall, the analysis of our Tcon/Treg cell epigenomic compendium, in conjunction with the T cell Hi-C data, implicated TFs associated with short-range looping and ultra-long-range metalooping in the T cell genome.

**Figure 7:**
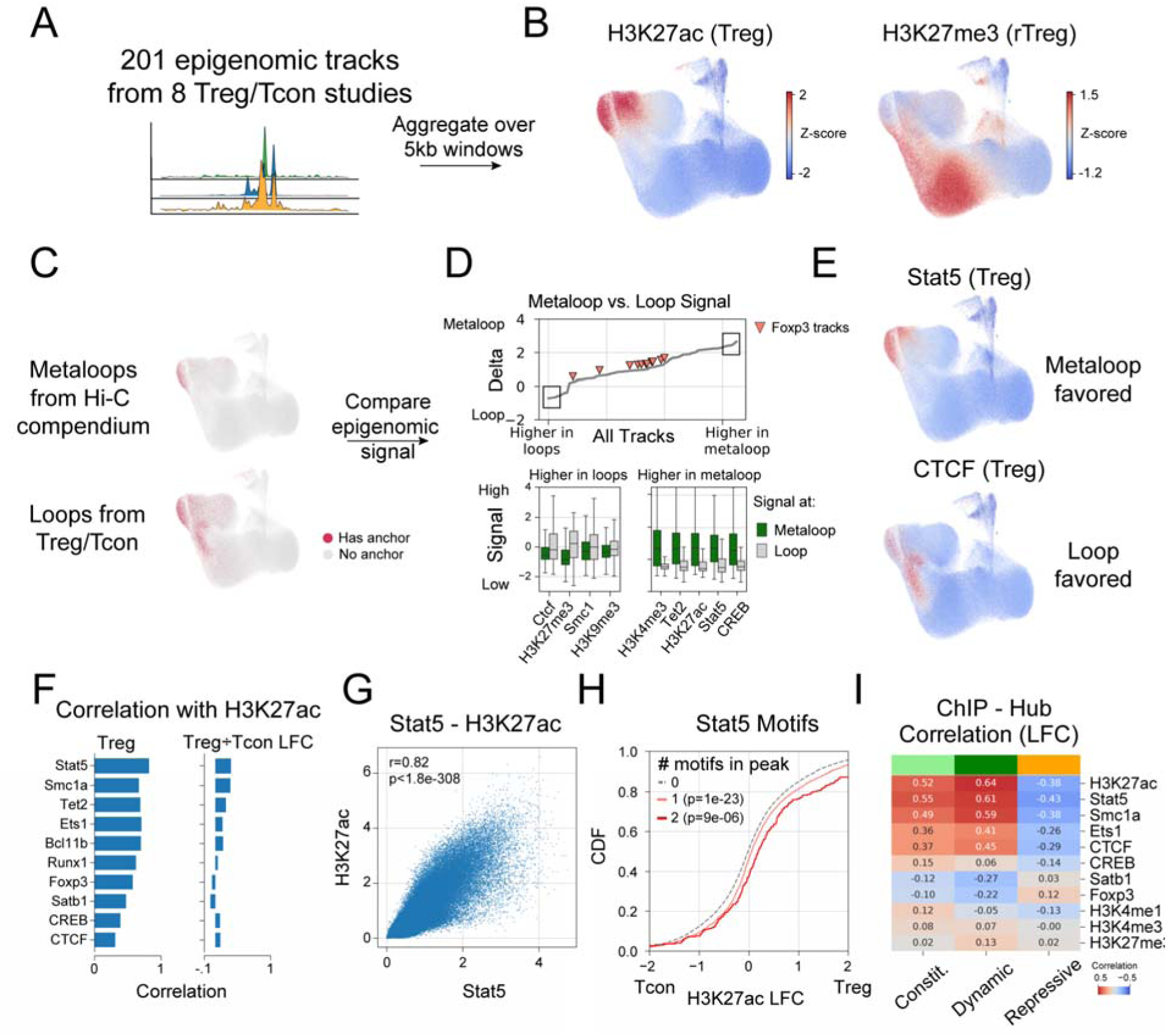
Integrative analysis of ChIP-seq, CUT&RUN and ATAC-seq data implicates metaloop-associated regulatory elements and transcription factors in Tcon and Treg cells. (A) Schematic of the approach. An epigenomic compendium of 201 ChIP-seq, CUT&RUN and ATAC-seq datasets was collected from eight Tcon and Treg cell studies. Each dataset was aggregated over 5 Kb bins tiling the genome and then z-score normalized. (B) Uniform Manifold Approximation and Projection (UMAP) embedding of the Tcon/Treg epigenomic compendium, where each point represents a 5 Kb genomic bin. Normalized Treg H3K27ac (left) and H3K27me3 (bottom) ChIP-seq signals are shown. (C) UMAP visualization of 5 Kb anchors of long-range metaloops (distance > 2 Mb) identified in Treg and Tcon metadomains (top), and anchors of short-range chromatin loops (< 2 Mb, bottom). These sets of genomic bins were used for differential epigenomic analysis. (D) Differential epigenomic analysis comparing long-range vs. short-range loop anchors (as in panel C). (Top) All factors ranked by enrichment at long-range vs. short-range loops, calculated as the difference in median ChIP-seq, CUT&RUN or ATAC-seq signal. (Bottom) Boxplots showing the most enriched factors at short-range (left) and long-range (right) interactions. Boxplots: center line, median; box limits, upper and lower quartiles; whiskers, 1.5x interquartile range. (E) UMAP visualizations of CTCF and Stat5 ChIP-seq signals in Treg cells. (F) Correlation between H3K27ac and TF ChIP-seq signals in Treg cells (normalized signal in Treg, left; log(Treg / Tcon), right) over all 5 Kb genomic bins. For Foxp3, log(Treg / Tcon) was replaced by log(Treg) since Foxp3 is not expressed in Tcon cells. (G) Scatterplot comparing Stat5 and H3K27ac ChIP-seq signals in Treg cells across 5 Kb genomic bins. (H) Differential H3K27ac ChIP-seq signal between Treg and Tcon cells in the H3K27ac ChIP-seq peaks, stratified by the number of Stat5 motifs per peak. (I) Correlation between differential hub metadomain score (Treg / Tcon) and differential ChIP-seq LFC signal (Treg / Tcon) for TFs and histone marks from panel F.

## Discussion

In this study we uncovered new principles of the 3D genome organization in primary mammalian T cells by developing novel approaches to analysis of chromatin organization data. We identified long-range and inter-chromosomal metadomain interactions between genomic regions of the size 50-250 kilobases, encompassing one or several genes. Our analysis revealed three major interchromosomal hubs in T cells, each encompassing dozens of genes and characterized by unique histone modification patterns and gene expression profiles, including two distinct active hubs and one repressive hub. The Active Constitutive hub, enriched for the promoter-associated histone mark H3K4me3, contains genes that are broadly expressed across immune cells and other tissues. In contrast, the Active Dynamic hub, enriched for the active enhancer-associated marks H3K4me1 and H3K27ac, predominantly contains T cell-specific genes and progressively strengthens during thymic differentiation. These observations suggest two distinct modes of gene activation occurring in spatially segregated nuclear compartments, likely governed by different molecular mechanisms. The increased strength of the Active Dynamic hub in activated T cells further suggests a role in facilitating the rapid transcriptional activation characteristic of immune responses. The spatial clustering of genomic loci with shared epigenetic features may also contribute to transcriptional robustness, as proposed in recent theoretical models (*58*). The Repressive hub, characterized by the polycomb-associated mark H3K27me3, included genes with lower expression, indicating a specialized repressive regulatory role.

Our characterization, while consistent with prior definitions of transcriptional hubs and condensate-like nuclear structures (*38*, *61*), extends beyond conventional models of genome compartmentalization such as A/B compartments and subcompartments. In particular, we describe metadomain hubs as higher-order assemblies of metadomains, focal cell type-specific interactions between distal TADs, organized through punctate enhancer-promoter contacts rather than diffuse compartmental interactions. While the Active Constitutive hub appears broadly conserved across cell types and is associated with nuclear speckles and housekeeping gene activation, the Active Dynamic hub is highly T cell-specific and has not been previously described. Furthermore, in previously defined subcompartments A1 and A2, the subcompartment A2 showed lower levels of activity with respect to a broad range of epigenetic marks, as indicated by lower levels of H3K4me1 and H3K4me3 and higher levels of H3K9me3 (*38*, *66*, *67*). In contrast, the Dynamic and Constitutive hubs show similar levels of activity, but distinct epigenomic profiles: the Dynamic hub is enriched for H3K4me1, while the Constitutive hub is enriched for H3K4me3. Notably, although previously reported H3K27me3-enriched subcompartment B1 was placed within the B compartment, we find that the Repressive hub, also marked by H3K27me3, remains largely within the A compartment. Nevertheless, further research is needed to better characterize physical properties and spatial positioning of the distinct nuclear structures associated with the metadomain hubs, e.g. through approaches such as TSA-seq, SPRITE, ORCA or seqFISH+ (*52*, *68*–*70*).

By aggregating many Hi-C datasets to achieve substantially increased resolution, we identified thousands of ultra-long-range metaloops occurring within metadomains. Metaloops connect 5 Kb genomic regions at distances of many megabases and frequently link promoters and enhancers. Using a straightforward extension of the ABC model of enhancer activity, we demonstrate that ultra-long-range and inter-chromosomal genomic interactions are significantly correlated with gene expression. Therefore, metaloops present an additional mode of gene regulation, distinct from classical short-range promoter-enhancer interactions and CTCF- and cohesin-mediated loops. Through integrative analysis of ChIP-seq, CUT&RUN and ATAC-seq data, we implicate several enhancer-associated TFs in ultra-long-range metalooping and confirm that CTCF and cohesin are depleted at anchors of these interactions. This is consistent with previous observations in cell culture models and neutrophils, where cohesin deactivation has been associated with emergence of long-range interactions (*11*, *14*). Future studies are needed to determine whether metaloops drive metadomain formation or whether metadomains facilitate specific long-range pairing of regulatory elements. Further dissection of metaloops, and identification of factors that distinguish subclasses such as Active Dynamic vs. Active Constitutive, may help explain how enhancer-promoter specificity is achieved across vastly different genomic distances. The analytical framework and algorithms we developed for Hi-C data analysis, applied both to individual datasets and at different levels of aggregation, will enable future discoveries of the regulatory principles underlying metaloops, metadomains and other features of chromosomal organization across scales and across biological systems.

## Acknowledgements

The authors thank the members of the Pritykin, Rudensky, Viny and Levine labs, and Christina Leslie, for helpful discussions. This work was supported by NIH grants DP2AI171161 (Y.P.), P30CA008748, R01 A1034206 (A.Y.R.), R01 AI138797, R01 AI153236, R01 AI146917, R01 AI168048, R01 AI107301, R01AI181664 and U19A171401 (all to A.P.), GM118147 (M.S.L.), R37 CA286857 (A.D.V). G.D. was supported by the NIH training grant T32 HG003284. A.N.N. was supported by the NIH/NIGMS training grant T32GM007388. X.H. was supported by the Cancer Research Institute Postdoctoral Fellowship. A.Y.R. is a Howard Hughes Medical Institute investigator. Y.P. was supported by the Ludwig Institute for Cancer Research and Princeton Precision Health. A.D.V. was supported by a Clinical Investigator grant from the Damon Runyon Cancer Research Foundation (120-22), a Clinician Scientist Development grant from the Doris Duke Charitable Foundation, and grants from Columbia University Vagelos College of Physicians & Surgeons (Gerstner Scholar Merit Award). A.P. was supported by Open Philanthropy, the Princeton Catalysis Initiative and Princeton University. We thank Christina DeCoste and the Molecular Biology Flow Cytometry Resource Facility, which is partially supported by the Rutgers Cancer Institute of New Jersey NCI-CCSG P30CA072720-5921 and an NIH S10 Shared Instrumentation Grant S10OD028592.

## Author contribution

Conceptualization: G.D., M.S.L., A.D.V., A.Y.R., and Y.P.; Methodology: G.D., Z.M.W., X.H., X.Y.B., W.K., A.D.V., A.Y.R., and Y.P.; Investigation: G.D., Z.M.W., X.H., S.S., M.J.W., X.Y.B., W.K., T.C., A.N.N., S.F., and A.D.V.; Formal Analysis: G.D., S.S., and Y.P.; Software: G.D., S.S., and M.J.W.; Data Curation: G.D.; Writing - Original Draft: G.D. and Y.P.; Writing - Review & Editing: G.D., S.S., M.J.W., A.P., P.S., M.S.L., A.D.V., A.Y.R., and Y.P.; Funding Acquisition: A.P., P.S., M.S.L., A.D.V., A.Y.R., and Y.P.; Supervision: A.P., P.S., M.S.L., A.D.V., A.Y.R., and Y.P.

## Data and code availability

The code used for analysis in this manuscript is available at https://github.com/pritykinlab/metadomain_paper. The metadomain calling algorithm and associated analysis pipeline is available at https://github.com/pritykinlab/InterDomain. Treg/Tcon, including aTreg/aTcon Hi-C data, and CD4 and CD8 T cell Hi-C data will be available at NCBI GEO upon journal publication. Results of reanalysis of published data are available in supplementary tables and alongside the code on github and the associated google drive folder.

## Methods

### Cell isolation

#### Treg/Tcon

A cell suspension was made from pooled secondary lymphoid organs (spleen, pLNs and mLNs) of Foxp3loxP-Thy-1.1-STOP-loxP-GFP (*Foxp3^GFP/LSL^*) mice (*40*, *41*) by meshing the organs through a 100-μm strainer (Corning, 07-201-432) with a syringe plunger. CD4+ T cells were enriched with the Dynabeads Flowcomp Mouse CD4 Kit (Thermo Fisher, 11461D) according to the manufacturer’s instructions, stained with antibodies, washed extensively, resuspended in isolation buffer (PBS with 2% FBS, 10 mM HEPES buffer, 1% l-glutamine and 2 mM EDTA), and sorted on a FACSAria (BD) instrument. Treg cells were sorted as TCRβ+CD4+CD8−NK1.1−Foxp3-GFP+Thy-1.1−, and naïve conventional CD4+ T cells as TCRβ+CD4+CD8−NK1.1−Foxp3-GFP−Thy-1.1−CD44loCD62Lhi.

#### Resting/Active Treg/Tcon/LSL

A cell suspension was made from pooled secondary lymphoid organs (spleen, pLNs and mLNs) of female Foxp3*^GFP/LSL^*and Foxp3*^DTR^* mice (*40*, *41*) by meshing the organs through a 100-μm strainer (Corning, 07-201-432) with a syringe plunger. CD4+ T cells were enriched with the Dynabeads Flowcomp Mouse CD4 Kit (Thermo Fisher, 11461D) according to the manufacturer’s instructions, stained with antibodies, washed extensively, resuspended in isolation buffer (PBS with 2% FBS, 10 mM HEPES buffer, 1% l-glutamine and 2 mM EDTA), and sorted on a FACSAria (BD) instrument. Wannabe Treg cells were sorted from Foxp3*^GFP/LSL^* mice as follows: naïve wannabe Treg cells: TCRβ+CD4+CD8−NK1.1−Foxp3-GFP−Thy-1.1+ CD44loCD62Lhi; activated wannabe Treg cells: TCRβ+CD4+CD8−NK1.1−Foxp3-GFP−Thy-1.1+CD44hiCD62L−; Treg cells and conventional CD4+ T cells were sorted from Foxp3*^DTR^* mice as follows: naïve Treg cells: TCRβ+CD4+CD8−NK1.1−Foxp3-DTR+ CD44loCD62Lhi; activated Treg cells: TCRβ+CD4+CD8−NK1.1−Foxp3-DTR+ CD44hiCD62L−; naïve conventional CD4+ T cells: TCRβ+CD4+CD8−NK1.1−Foxp3-DTR−CD44loCD62Lhi; activated conventional CD4+ T cells: TCRβ+CD4+CD8−NK1.1−Foxp3-DTR− CD44hiCD62L−. 100,000 cells were sorted for each population per replicate.

All studies were approved by the MSKCC Institutional Animal Care and Use Committee under the protocol 08-10-023.

#### CD4 and CD8 T cells

Mice: 6-week-old female C57BL/6J mice were obtained from The Jackson Laboratory (Bar Harbor, ME). Experiments using C57BL/6J mice were performed in accordance with a protocol (number 3063), which was reviewed and approved by the Institutional Animal Care and Use Committee (IACUC) of Princeton University.

Isolation and preparation of CD4 and CD8 T cells from mouse spleen: Mice were euthanized by intraperitoneal injection of ketamine/xylazine at a supratherapeutic dose. Spleens were collected and placed into 5 mL of serum-free Dulbecco’s Modified Eagle Media (DMEM, Thermo Fisher Scientific, Waltham, MA, 11995081). Individual spleens were placed on 100-μm strainers then mechanically dissociated using the plunger of a 3 mL syringe. The strainer was washed with 5 mL of serum-free DMEM in which the splenocytes were subsequently resuspended. The single cell suspension was centrifuged at 1,300 rpm for 5 min at 4°C, then the pellet was resuspended in 1x BD Pharm Lyse lysis buffer (BD Biosciences, Franklin Lakes, NJ, 55599) and incubated at room temperature in the dark for 15 min. Lysis was quenched using DMEM supplemented with 10% (v/v) FBS and 1% (v/v) penicillin-streptomycin. The splenocyte suspensions were spun at 1,300 rpm for 5 min at 4°C, then washed with FACS buffer (1% (v/v) FBS in PBS). The cells were counted using 0.4% trypan blue solution (Thermo Fisher Scientific, 15250061) and a hemocytometer, then resuspended at a concentration of 1 x 107 cells/mL. An aliquot of cells was set aside as an unstained control. Another aliquot of cells was set aside to generate the live/dead control in which half of the cells are heat shocked at 95°C for 5 min, set on ice for 2 min, and combined with the live cells.

Antibody staining and fluorescence-activated cell sorting: The following fluorophore-conjugated antibodies were used for flow cytometry: CD45-FITC (clone 30-F11, 1.25 μg/mL, BioLegend, San Diego, CA 103108), CD3-Alexa700 (clone 17A2, 1 μg/mL, Invitrogen, Waltham, MA, 56-0032-82), CD4-PE-Cy7 (clone RM4-5, 1 μg/mL, Invitrogen, 25-0042-82), CD8-BV510 (clone 53-6.7, 1 μg/mL, BioLegend, 100752). The antibody cocktail was prepared in FACS buffer, then incubated with cells for 30 min at 4°C in the dark. The splenocyte suspensions were washed 2x in FACS buffer. To serve as a viability marker, DAPI (1 μg /mL, Sigma-Aldrich, St. Louis, MO, D9542) was directly added to antibody-stained cells and the live/dead control 15 min prior to FACS acquisition. Compensation controls were prepared using AbC Total Antibody Compensation Bead Kit (Invitrogen, A10497). The FACSAria Fusion Flow Cytometer (BD Biosciences) or the MA-900 cell sorter (Sony Biotechnology, San Jose, CA) was used to perform cell sorting and acquired data were analyzed using BD FACSDiva or MA900 software. For collection of each T cell subset, the samples were gated as follows: for CD3+ T cells, CD45+ CD3+; for CD4+ T cells, CD45+ CD3+ CD4+ CD8- ; and for CD8+ T cells, CD45+ CD3+ CD4-CD8+. Cells were sorted into FACS buffer and maintained at 4°C.

#### Hi-C for Tcon/Treg and CD4/CD8 T cells

Hi-C was performed as previously described (*71*). Briefly, sorted T cell populations (approximately 1×10^5^) were cross-linked in 1% formaldehyde for 10 minutes and quenched in 125mM glycine. Cross-linked cells were lysed and chromatin was digested using the Arima HiC^+^ kit (Arima Genomics, San Diego, CA) using protocol adaptations for low cell input. Digested and reverse crosslinked DNA was eluted in 100uL and fragmented to 350 bps using a Covaris LE220Rsc sonicator (Covaris, Woburn, MA). Sheared genomic material was enriched for biotinylated DNA using streptavidin beads followed by library preparation using Arima protocol modifications for Accel-NGS 2S DNA plus library kit (IDT, Coralville, IA). After end repair and ligation, libraries were quantified using the KAPA library quantification kit (Roche, Indianapolis, IN) and PCR amplified for the number of cycles required to generate >4nM per library. Hi-C libraries were sequenced on an Illumina NovaSeq and raw sequencing data in the FASTQ format were obtained.

#### Hi-C data preprocessing

Hi-C reads were aligned to the mouse UCSC mm10 GRCm38 genome (chromosomes 1-19, X, Y) downloaded from the 4DNucleome (4DNFI2493SDN) using bwa mem version 0.7.17, with settings --SP5M. Pairs files were generated using pairtools parse version 0.3.0 with a min-mapq cutoff of 30 and --walks-policy 5unique, followed by pairtools dedup with default parameters, and pairtools select for the following pair types: UU, UR, and UR. Pairs files were mapped into cool files using cooler version 0.8.11. Hi-C data in these cool files was normalized using cooler (*72*) balance with default parameters for each sample separately and for merged biological replicates of each condition across all conditions. This way, balancing normalization was applied to both intrachromosomal and interchromosomal Hi-C signal.

#### Loop and TAD calling

Loops were called using Mustache (*73*) version 0.1.9 at 5 Kb resolution with the following parameters: r = 5,000, FDR = .02, st = .85, sz = 1.2. Loops were called in three different ways to maximize the number of loop calls: (1) merged Treg replicates, (2) merged Tcon replicates, and (3) merged Treg and Tcon replicates. Loops from these three calls were merged into a single loop atlas (n=18,098). To avoid redundancy, we selected only a single representative among the loops from the three loop calls which were ≤ 2 bins in distance at both anchors. In other words, for example if a loop from Treg or Tcon cells was close to a loop from the merged Tcon/Treg data, only the loop from the merged data was retained. This was done iteratively in the following order: Merged > Treg > Tcon. This merging procedure yielded 13,225 loops. TADs were called using cooltools (*74*) version 0.5.4 using the diamond-insulation function, with window_bp=16. Loops crossing TAD boundaries were defined using Bedtools pairToBed with type=’ispan’, so that a loop crossing a TAD boundary was only defined if the span between (but not including) the loop anchors overlapped one or more TAD boundaries.

#### Differential looping analysis

Using the loop atlas described above, we obtained read counts for each Tcon and Treg replicate using Cooler. DESeq2 (*75*) was used to identify loops with statistically significant differential read counts between Treg and Tcon cells. To account for both sample-specific sequencing depth and distance decay, we used scaling factors calculated separately for loops of different genomic distances. Specifically, for each chromosome, we binned loops into 5 groups, equidistant on a log scale based on genomic distance between loop anchors (a max length of 2Mb), and calculated separate size factors for all loops in each group using DESeq2 estimateSizeFactors(). Then DESeq2 was called jointly for all groups. Loops with |LFC| > 0 and FDR < .05 were called significantly differential, yielding 1,679 Tcon-specific loops and 1,008 Treg-specific loops (LFC; log2 fold change). Treg-specific loop anchors were defined as all anchors of Treg-specific loops, i.e. with significantly higher Treg Hi-C reads, and similarly for Tcon-specific loop anchors. This yielded 3,023 Tcon-specific loop anchors, 1,839 Treg-specific loop anchors, 66 genomic regions that were anchors of both Tcon- and Treg-specific loops, and 15,945 genomic regions that were anchors of loops that were not significantly differential between Tcon and Treg.

#### Hi-C and other epigenomic data visualization

Balanced Hi-C data was plotted using Coolbox (*76*). Gene annotation plotting was performed using pygbrowse (*77*) for GTF annotations from Ensembl version GRCm38.93. ChIP-seq data were preprocessed and normalized using MACS2 (*78*), and bigWig files produced (see section below for details). BigWig tracks were plotted with custom code (see Github repository).

#### Gene expression analysis

RNA-seq data (read count matrices) for resting Tcon and resting Treg cells were obtained from NCBI GEO (accession number GSE154680). Differential gene expression analysis was performed using DESeq2. Genes with |LFC| > .25 and FDR < .05 were called significantly differentially expressed. Protein-coding genes with an average read count of >= 4 were kept for subsequent analysis. This yielded 12,527 genes, of which 1,583 were significantly overexpressed in Tcon and 554 were significantly overexpressed in Treg cells. RNA-seq RPKM (reads per kilobase million) values were calculated by multiplying all read counts by the same factor so that they sum to 1e6 and dividing by gene lengths (in Kb). Gene lengths were calculated using the FeatureCounts ‘Length’ column.

#### Comparison of differential gene expression and differential looping

For each gene, we considered all loops with an anchor within 10 Kb of the gene TSS, defined as the start of the transcript with the highest support level in annotations from Gencode mouse release M23. Then, all DESeq2-estimated Hi-C read count LFC values for those loops were averaged for each gene. Genes which did not intersect any loop anchor were not considered for this analysis.

#### ABC score analysis

Activity-by-contact (ABC) score for a TSS and a distal region *R* on the same chromosome was calculated as the product between MACS2-normalized H3K27ac ChIP-seq signal at *R* and the balanced Hi-C signal for the interaction between the TSS and *R*, both binned at 5 Kb resolution (ChIP-seq bigWig averaged over a bin). The analysis was performed separately in Tcon and in Treg cells. ABC values for genomic regions within 10 Kb from a TSS were set to zero, to avoid local confounding effects. To calculate the cumulative ABC score for a single TSS, we summed all ABC scores for genomic regions within a certain distance from the TSS. We considered the distance cutoffs of 50 Kb, 100 Kb, 1 Mb, 10 Mb, and without the cutoff, i.e. for the entire chromosome. As a control, we performed the same calculation, but with two modifications. To control for contribution from the H3K27ac ChIP-seq signal, we shifted H3K27ac ChIP-seq signal by 200 Kb (and with intact actual Hi-C signal). To control for contribution from the Hi-C signal, we randomly shuffled intrachromosomal Hi-C along each diagonal within each chromosome. For interchromosomal Hi-C, we shuffled all values within each bin. For each chromosome, for all genes expressed in Tcon or Treg cells, we calculated Pearson correlation between RNA-seq RPKM for a gene and the cumulative ABC score for the TSS of that gene, both log-transformed. This correlative analysis was done separately for Tcon and Treg cells. We also calculated Pearson correlation between RNA-seq LFC values (between Treg and Tcon) for a gene and the LFC in cumulative ABC scores (between Treg and Tcon) for the TSS of that gene. These calculations were done for the cumulative ABC scores defined using different distance cutoffs. We also generalized definitions of ABC and cumulative ABC to interchromosomal analysis in a straightforward manner, applying it to balanced interchromosomal Hi-C signal.

#### Observed/expected (O/E) matrix

The intrachromosomal Hi-C observed vs. expected (O/E) matrix was calculated by taking the balanced Hi-C matrix and dividing each diagonal (set of matrix elements corresponding to interactions at the same fixed genomic distance) by the average Hi-C value along that diagonal. This was done for each chromosome separately. A pseudocount of 1e-4 was added to both the observed and the expected matrices to account for sparsity. Lastly, the logarithm of the O/E was taken to compute the Log(O/E) matrix. For interchromosomal contacts, the balanced Hi-C matrix corresponding to each pair of chromosomes was divided by the average contact, and the logarithm was taken. A pseudocount of 1e-4 was added to both the observed and the expected matrices to account for sparsity. O/E was calculated at 250 Kb, 50 Kb and 25 Kb resolution.

#### A/B compartment analysis

Intrachromosomal A/B compartment score was called as previously described in (*49*). First, the pairwise Pearson correlation matrix of the Log(O/E) matrix was calculated. Then, we calculated the PCA decomposition of the Pearson correlation matrix using sklearn version 1.3.0, and the compartment score was set to the first principal component. The sign of the compartment score was aligned with the A compartment using H3K27ac ChIP-seq signal. To make compartment scores comparable between chromosomes, compartment vectors were normalized to have an L2 norm of N, where N is the number of bins in that chromosome. Compartments were calculated at both 50 Kb and 250 Kb resolutions.

#### Chromosome-wide differential Hi-C analysis over 250 Kb bins

We aggregated read counts for each replicate of each biological condition at 250 Kb resolution for each chromosome separately. DESeq2 was used to identify interactions between 250 Kb genomic regions with statistically significant differential Hi-C read count between Treg and Tcon cells. Interactions with an average read count of less than two were discarded. To account for both sample-specific sequencing depth and distance decay, we used scaling factors calculated separately for interactions of different genomic distances. Specifically, for each chromosome, we binned interactions into 50 groups, equidistant on a log scale based on genomic distance between 250 Kb interaction anchors, and calculated separate size factors for all interactions in each group using DESeq2 estimateSizeFactors(). Differential Hi-C was then calculated using DESeq2 with cutoffs |LFC| > 0 and FDR < .05. This yielded 114546 Treg-specific interactions and 91415 Tcon-specific interactions. Wald statistic for differential Hi-C signal accounting for both the magnitude and variance, as calculated by DESeq2 in this analysis, was used for visualization.

#### InterDoman algorithm for metadomain calling

The new algorithm InterDomain identifies metadomains genome-wide. It was inspired by the HICCUPS loop calling algorithm (*38*). Metadomains were defined as interactions between large genomic regions (e.g. of length 250 Kb) that have statistical read count enrichment and prominence. For this, InterDomain was applied to normalized intrachromosomal Hi-C data at coarse resolution, e.g. 50 Kb. First, InterDomain identifies prominent pixels in the 2D Log(O/E) matrix. Prominent pixels are local maxima whose topographic prominence in both the X and the Y direction exceeds a cutoff of 4. Topographic prominence is defined as the lowest drop in height along the path from one local maximum to the nearest larger local maximum. After identifying prominent pixels, InterDomain computes read count enrichment at each prominent pixel relative to a neighborhood of nearby pixels serving as the local control. InterDomain assumes that reads are drawn from a Poisson distribution parameterized by the average signal of the local control. Prominent peaks with a read count enrichment p-value < 1e-20 are designated as metadomains. Lastly, to prevent double-counting and redundancy, all immediately adjacent pixels which are called as metadomains are collapsed into one call where the pixel with the greatest Log(O/E) signal is selected as a representative of the metadomain. For interchromosomal analysis, InterDomain is run with the following modifications. Identification of prominent pixels is done on a smoothed O/E matrix. Smoothing was performed using scipy.ndimage.gaussian_filter with sigma=.75. This is done to overcome the sparsity and lower read count of the interchromosomal matrix. Then read count enrichment is calculated using a pseudocount of .5 to prevent high sensitivity in highly sparse regions. Finally significant metadomain calls are identified using a less stringent p-value cutoff of 1e-5, due to the lower statistical power from lower read counts. For both inter- and intrachromosomal analysis, InterDomain was run on 50 Kb resolution Tcon and Treg cell Hi-C data to generate metadomain calls. These 50 Kb metadomains were then analyzed at 250 Kb resolution by overlapping them with the 250 Kb bins that had been used for Hi-C analysis.

#### Intrachromosomal metadomain analysis

For metadomain triplet enrichment analysis, a metadomain triplet was defined as three 250 Kb bins in which each pair of bins is connected by a metadomain. For each chromosome, triplet calculation was performed on the intrachromosomal metadomain matrix as well as a randomly shuffled matrix as a control. 500 permutations were performed to determine statistical significance. For comparison of metadomains between mouse and human, genomic regions of length 50 Kb (corresponding to the original 50 Kb metadomain calls) in mouse genome mm10 were lifted over to human genome hg38 using the online web tool UCSC LiftOver with the following procedure. For n=52,685 50 Kb metadomain anchors, 121,731 corresponding regions were identified using LiftOver due to individual regions being split. To disambiguate, we first discarded results in hg38 that were less than 10 Kb or greater than 200 Kb. Second, for bins which lifted over to multiple genomic locations, we chose the genomic location with the largest length in human as the unique representative of this bin. This allowed mapping of the mouse Tcon and Treg metadomain 50 Kb bins to human. Out of 52,685 metadomain bins, 45,451 lifted over (including intra-chromosomal and inter-chromosomal). For analysis of intra-chromosomal metadomains, we required that both metadomain anchors map over successfully, and to the same chromosome. For analysis of inter-chromosomal metadomains, we only required that both metadomain anchors map over successfully. For analysis of superenhancers (SEs) in metadomains, SEs were obtained from (*28*) and then lifted over from mm9 to mm10 using UCSC LiftOver. For analysis and visualization of ChIP-seq signal in metadomains, MACS2-normalized ChIP-seq signal was aggregated over 250 Kb genomic bins. Specifically, ChIP-seq signal was averaged over 250 Kb bins using the Python package pybbi (v0.3.2) and then log-transformed. Intrachromosomal metadomains were clustered for each chromosome separately using hierarchical clustering from scipy (scipy.cluster.hierarchy) (v1.11.2) with method=’ward’ and metric=’euclidean’ and n_clusters=10.

#### Genome-wide metadomain clustering and interchromosomal metadomain hubs

First, inter- and intra-chromosomal metadomains were combined across Treg and Tcon to create a matrix representing all detected metadomains. Second, bins which had more than either at least 20 inter-chromosomal or at least 20 intra-chromosomal metadomains were included. Third, the metadomain matrix was transformed using sklearn PCA version 1.3.0 with 20 components. Scipy’s hierarchical clustering with method=’ward’ and metric=’euclidean’ was applied to the PCA loadings of each bin to generate 24 unique clusters. Clusters with a high concentration of just one chromosome were removed and clusters with low (< 5%) metadomain density (defined as the fraction of all interchromosomal pairs containing a metadomain) were removed, producing a final list of metadomain clusters. Genome-wide metadomain clustering identified very fine-grained metadomain clusters. To aggregate these results and merge similar clusters, we can calculate the average number of metadomains between clusters, creating an adjacency matrix between clusters. Louvain clustering algorithm (python-louvain v0.16) can be run on this adjacency matrix to define groups of metadomain clusters sharing many metadomains. For our Tcon and Treg Hi-C data, we performed this analysis and ran the Louvain clustering at resolution 1 and thus defined three groups of metadomain clusters that formed three interchromosomal hubs.

#### Comparison of metadomain clustering with Hi-C clustering

To confirm the robustness of clusters formed by metadomain clustering, we formed a matrix of the same bins using Log(O/E) Hi-C data as the input, instead of binary metadomain as input. For this matrix, we ran an identical clustering procedure to the one that was run on our metadomains and compared the resulting clusters with metadomain clusters.

#### ImmGen data analysis

We reanalyzed RNA-seq gene expression data for different immune cells from the ImmGen consortium (*39*). Processed RNA-seq read counts were obtained from NCBI GEO (accession number GSE109125). RPKM values were calculated and Z-scored across all genes within each cell type. Z-scores were then averaged for all genes overlapping each metadomain hub or an equivalently strong set of bins in the A compartment (distinct from bins in hubs) to generate “hub activity” scores. Hub activity scores were clustered using hierarchical clustering.

#### Mouse sci-ATAC-seq atlas data analysis

Processed ATAC-seq matrices were downloaded from (*51*). ATAC-seq signal was pseudo-bulked by annotated cell type and peaks were z-score normalized across cell types. Peaks overlapping hub annotations were averaged and compared across cell types and different hubs.

### Housekeeping gene data analysis

A list of housekeeping genes were downloaded from (*79*) and the fraction of housekeeping genes in each hub was calculated.

#### Functional term enrichment analysis

Functional enrichment analysis was performed by taking all genes overlapping all bins within a metadomain hub. These genes were passed into the Python package gprofiler v1.0.0 with the following sources: GO MF, GO BP, GO CC, KEGG, REAC, WP. All genes passing previously described filtering steps were used as a control.

#### Metadomain score calculation (hub level)

To calculate metadomain scores for an entire hub, we generated interchromosomal pileups at 50 Kb or 25 Kb resolution between groups of bins, such as the Active Constitutive or Active Dynamic hubs. These pileups were run on the Log(O/E) matrices. Specifically, to calculate the Metadomain Score, we calculated the average Log(O/E) signal between all interchromosomal pairs of bins in a hub (“inside” values), and subtracted the average Log(O/E) signal of flanking regions (“outside” values). A Mann-Whitney U test between inside and outside values was run to compute a p-value.

#### Metadomain score calculation (individual bin)

To calculate metadomain scores for an individual bin of interest, we followed the same protocol as above. Specifically, to calculate the Metadomain Score (MS), we calculated the average Log(O/E) signal between that bin and all bins in the hub from a different chromosome than the bin of interest (“inside” values). We then subtracted the average Log(O/E) signal of regions flanking the bin of interest (“outside” values). A Mann-Whitney U test between inside and outside values was run to compute a p-value.

#### Analysis of SE metadomain scores

Metadomain scores were calculated for each SE and the Active Constitutive and Active Dynamic hubs. SEs with a MS p-value less than 1e-20 and a MS score greater than .03 with either of the Active hubs were designated as having a recovered “pileup” metadomain and included in the plot in Figure 4F.

#### Differential metadomain scores

To determine differential metadomain scores, we calculated metadomain scores in each condition (Treg/Tcon) separately and calculated the difference between the two metadomain scores. A p-value was calculated using a Mann-Whitney U test comparing inside-outside enrichment between the two conditions. This calculation was performed both at the hub level (i.e. all pairs of contacts in a hub) and for bin-hub contacts (i.e. all contacts between one bin and the hub).

#### Single-cell RNA-seq co-expression analysis for metadomains and metadomain hubs

Single-cell RNA-seq data for Treg cells (GSM3978655) and their precursors (GSM3978654) was obtained from NCBI GEO (accession number GSE134902). The data was read into Scanpy (v1.9.4), aggregated into metacells (n=72, precursors; n=69, Treg) using Leiden clustering to reduce sparsity, and normalized using Pearson residual normalization. Then, Pearson correlations were calculated for all pairs of genes. We examined enrichment in correlations between genes in metadomains (“baseline co-expression”), as well as Treg-specific enrichment in correlations for genes that have Treg-specific metadomains (“differential co-expression). To determine baseline co-expression, we took correlations between pairs of genes where both genes were in a metadomain hub and compared this with correlations between pairs of genes where only one gene was in a metadomain hub. Mann-Whitney U test was used for statistical comparison. For differential co-expression, we examined all genes which were not originally in the hub but which displayed Treg-specific hub contact (“hub-up” genes). We then took correlations in Treg cells between hub-up genes and other genes in the hub and compared this with correlations between hub-up genes and non-hub genes. We performed the same comparisons in precursor cells as a control.

#### Graph embedding of differential metadomains in a metadomain hub

To generate a graphical embedding of a hub based on differential metadomaining patterns, we subtracted the Treg metadomain hubs from the Tcon metadomain hubs to create a matrix representing differential metadomaining. Then, we computed pairwise Pearson correlations of the differential metadomain matrix. A high correlation means that two bins have similar differential metadomains, whereas a negative correlation means that the two bins have opposite differential metadomains. To prevent numerical instability in the graph layout, negative correlations were clipped to -.00001 while positive correlations were clipped at .5, and self-correlation was set to zero. This matrix was used as input to the NetworkX version 3.1 spring layout embedding algorithm, which was run with default parameters to calculate a 2D graph spring layout corresponding to differential metadomaining patterns. The 2D graph hub embedding was visualized with the following settings. Nodes, representing each bin, had a size that was proportional to the number of Treg-specific or Tcon-specific metadomains for that bin (whichever was larger). Nodes were connected by edges if the two bins were connected by a differential metadomain. Edge color was blue for Tcon-specific metadomains and red for Treg-specific metadomains. Shared metadomains were not plotted as edges. Bins with fewer than 25 Treg-specific or Tcon-specific metadomains with the hub were pruned and their edges were removed.

#### T cell Hi-C compendium

Hi-C datasets generated in T cells were identified using SRA Explorer by searching for terms such as Hi-C, HiC, T-cell, and Tcell. In total, we identified 60 datasets from 17 studies (Table S4). Hi-C was aligned and processed as described above. To identify Hi-C datasets with similarity to our Treg/Tcon Hi-C data, we analyzed all Hi-C datasets at a resolution of 250 Kb and calculated O/E matrices for each dataset. Datasets with a correlation greater than .7 with the Treg Hi-C dataset were selected, resulting in 51 datasets from 15 studies. The selected datasets were merged to form a “mega” Hi-C dataset that contained 19 billion processed read pair contacts. This dataset was normalized through balancing as described above. A comprehensive list of included Hi-C datasets can be found in Table S4.

#### Metaloop calling algorithm

A modified version of InterDomain was used to refine 50 Kb metadomains identified in Treg or Tcon cells by identifying 5 Kb focal interactions in the T cell Hi-C compendium dataset. Metadomain refinement was performed only at metadomains that had been identified at 50 Kb resolution in the Treg or Tcon data. For each 50 Kb metadomain, Hi-C data at the metadomain and in a 100 Kb radius was analyzed at 5 Kb resolution in the T cell compendium dataset. Intrachromosomal InterDomain was then run on this data matrix with the following modifications. First, read count enrichment was thresholded with a p-value cutoff of 1e-15 instead of 1e-20. Second, unlike intrachromosomal InterDomain, we did not limit the analysis to “prominent peaks” since we had already restricted our analysis to previously called metadomains. The same process was run at metadomains shifted by 250 Kb in both anchors as a control.

#### Single-cell ATAC-seq co-accessibility analysis

Single-cell ATAC-seq data for splenic Treg cells (*80*) was obtained from NCBI GEO (accession number GSE156112; GSM4724883; GSM4724889). The count matrices were read into Scanpy (*81*) (v1.9.4). Metacells were formed to reduce sparsity according to the previously described protocol used for the Cicero algorithm (*82*). Pearson residuals were calculated on metacell data. Then, Pearson correlations were calculated for different pairs of 5 Kb bins.

#### ChIP-seq, ATAC-seq and CUT&RUN data analysis

SRA Explorer and a literature search were used to identify published ChIP-seq, ATAC-seq and CUT&RUN data for Tcon and Treg cells. In total, we identified 201 experiments from 8 studies (Table S1). Reads were aligned to the mouse genome mm10 using bowtie2 (*83*) (v2.5.0) with parameters --very-sensitive --no-unal -- no-mixed --no-discordant. For ChIP-seq and ATAC-seq, bigWig files were generated using macs2 callpeak and macs2 bdgcmp. For ChIP-seq, inputs were passed into MACS2 when available. For CUT&RUN, bigWig files were generated using deeptools bamCoverage (v3.5.2) with parameters -bs 10 --normalizeUsing RPGC, since MACS2 assumptions were not appropriate for normalization of CUT&RUN data. Only tracks profiling Treg cells, Tcon cells, Treg precursor cells, or Foxp3-GFPKO cells were included in the final analysis. ChIP-seq, ATAC-seq and CUT&RUN signal from the bigWig files generated above was averaged over each 5 Kb genomic bin and log1p-transformed. Then, Z-scores were calculated within each experiment across all bins. Then bins intersecting the ENCODE blacklist were removed (n=57,944; 11% of all 5 Kb bins) (*84*), and bins with low levels of signal across all sites (Z-score < 1.5 for every measured track, n=78,085; 16% of remaining 5 Kb bins) were excluded, leaving 487,666 bins. Scanpy (v1.9.4) was used to read in the Z-scored data, and a kNN graph was constructed in PCA space using 20 components, 30 neighbors, and cosine similarity as the metric. UMAP coordinates were generated using default Scanpy parameters. To calculate differential epigenomic signal (ChIP-seq, ATAC-seq or CUT&RUN) across two sets of genomic bins of interest (e.g. corresponding to peaks, loop anchors, or other annotations), we subset the epigenomic signal only to these bins, and then recalculated Z-scores of the epigenomic track along these subsets jointly. For calculation of epigenomic signal in metaloop anchors, we restricted to those metaloop anchors which overlapped short-range loop anchors. Then, we calculated the difference in Z-scores across the two sets of bins for each factor using a Mann-Whitney U test to calculate p-values. The difference of the median Z-score across each set of bins was used to rank factors.

#### Motif calling and enrichment analysis

Motifs were called using FIMO (v5.5.0) with the JASPAR2022 CORE non-redundant vertebrates motif database and a p-value cutoff of 5e-5. Motifs were called at H3K27ac ChIP-seq peaks (n=42179) or ATAC-seq peaks (n=93417) identified in Treg and Tcon cells using MACS2. For motif enrichment analysis in loop anchors, we subset analysis to motifs in ATAC-seq peaks overlapping loop anchors. To compare motif enrichment between different sets of loop anchors, we calculated the fold change in the fraction of ATAC-seq peaks with each motif, for ATAC-seq peaks between the different sets of anchors. For metaloop analysis, we restricted to metaloop anchors overlapping short-range loop anchors. To calculate p-values, we used a Fisher exact test for each motif; these were corrected for multiple hypothesis testing using the Benjamini-Hochberg procedure. For motif enrichment in hubs, we used all ATAC-seq peaks overlapping the 250 Kb metadomain hub bins. To identify motifs associated with differential H3K27ac between Treg and Tcon cells, we compared the H3K27ac ChIP-seq LFCs for peaks containing a motif compared to all other peaks. Specifically, we calculated the mean difference in H3K27ac LFC and calculated p-values using a Mann-Whitney U test.

#### MACS2 Peak calling for H3K27ac and ATAC-seq

Peaks were called in H3K27ac ChIP and ATAC-seq datasets by running macs2 callpeak on individual replicates with a p-value threshold of .01 and filtering to reproducible peaks (IDR < .1) (*85*). Then, reads mapped to ATAC-seq peaks were quantified using Rsubread (*86*) and DESeq2 was used to identify differentially accessible or acetylated peaks.

## Supplementary Tables

**Table S1:** Characteristics of new and published Hi-C data and published ChIP-seq data used in this study.

**Table S2:** Tcon and Treg Hi-C loops and TADs.

**Table S3:** Global differential Hi-C analysis at 250 Kb resolution in Tcon and Treg cells.

**Table S4:** Tcon and Treg Hi-C metadomains.

**Table S5:** Metaloops from the T cell Hi-C compendium.

