## Supplementary Figures for "Metadomain and metaloop genome interactions in mammalian T cells"

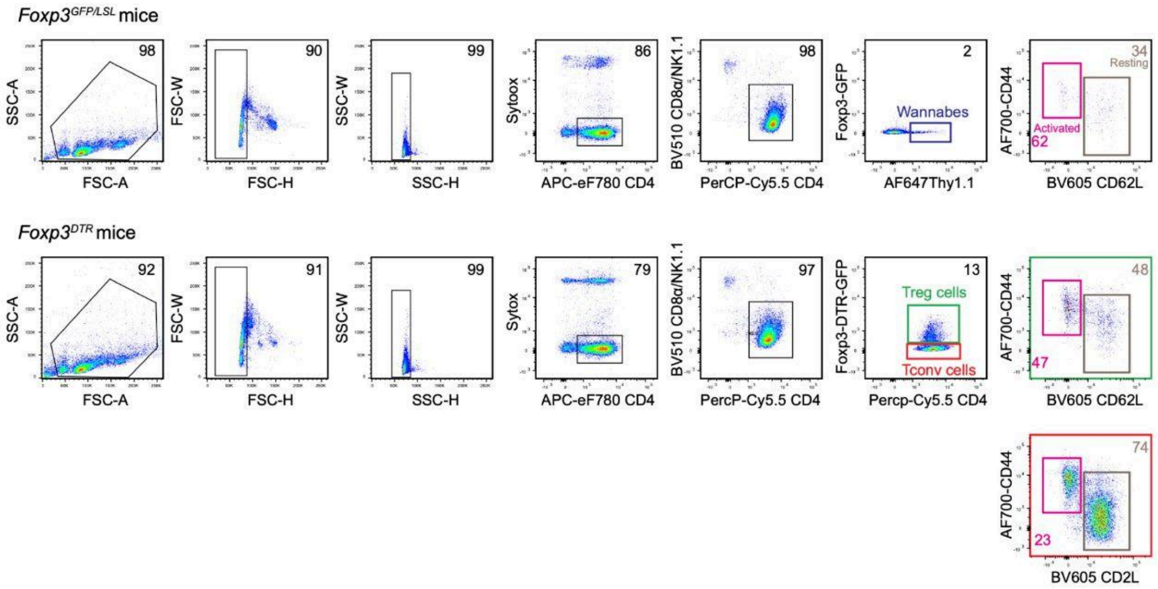


**Figure S1: Flow cytometry gating strategy. Related to Figures 1 and 5.** Treg and Tcon cells were isolated from Foxp3$^{GFP/LSL}$ mice. Activated and resting Treg and Tcon cells were isolated from Foxp3$^{DTR}$ mice. See Methods for details.

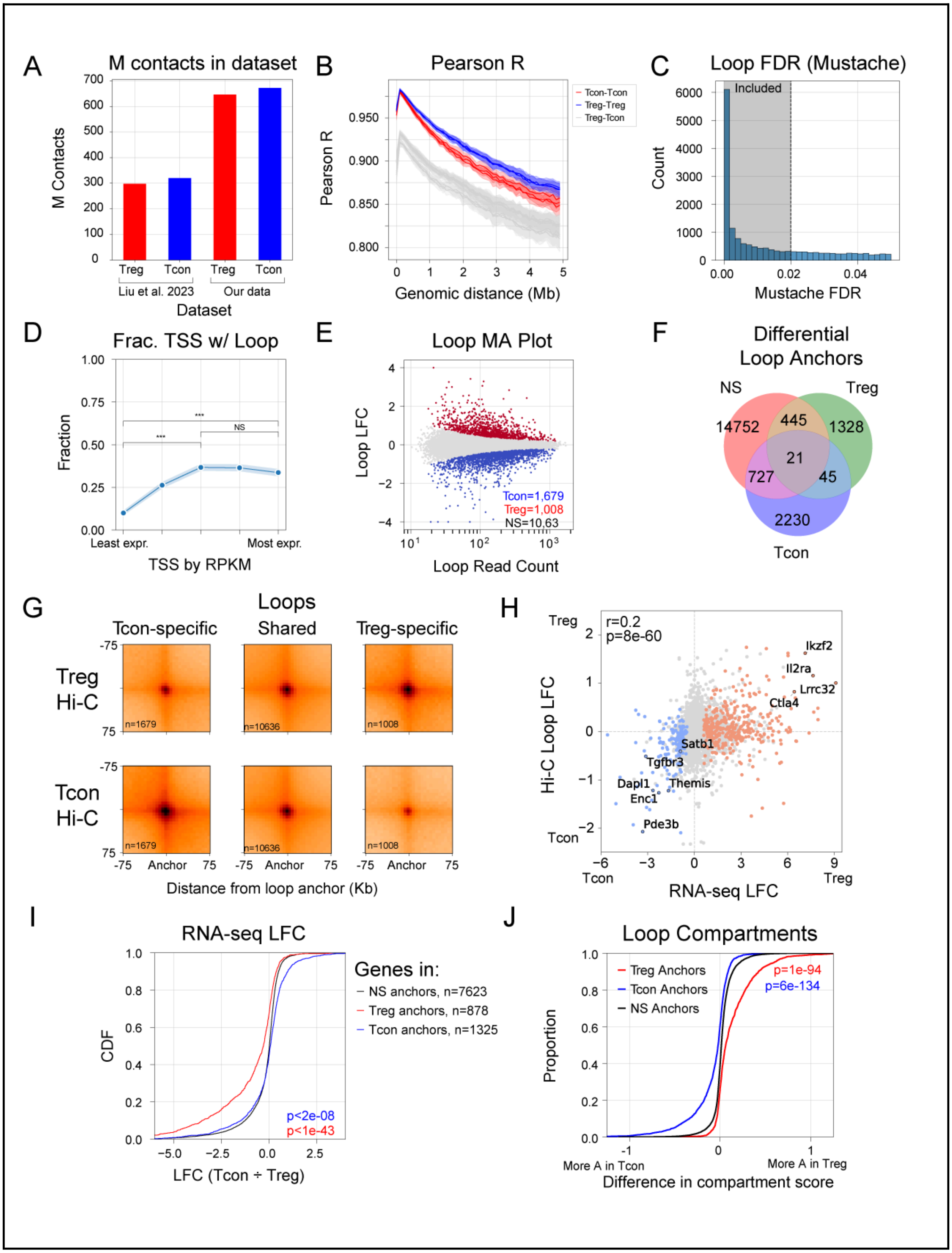
A
M contacts in dataset
M Contacts
Treg
Tcon
Treg
Tcon
Liu et al. 2023
Our data
Dataset
B
Pearson R
Tcon-Tcon
Treg-Treg
Treg-Tcon
Genomic distance (Mb)
C
Loop FDR (Mustache)
Included
Count
Mustache FDR
D
Frac. TSS w/ Loop
Fraction
Least expr.
Most expr.
TSS by RPKM
E
Loop MA Plot
Loop LFC
Tcon=1,679
Treg=1,008
NS=10,63
Loop Read Count
F
Differential
Loop Anchors
NS
Treg
Tcon
14752
445
1328
21
727
45
2230
G
Loops
Tcon-specific
Shared
Treg-specific
Treg
Hi-C
Tcon
Hi-C
n=1679
n=10636
n=1008
Anchor
Distance from loop anchor (Kb)
H
r=0.2
p=8e-60
Hi-C Loop LFC
Ikzf2
Il2ra
Lrrc32
Ctla4
Satb1
Tgfbr3
Themis
Dapl1
Enc1
Pde3b
RNA-seq LFC
I
RNA-seq LFC
CDF
Genes in:
NS anchors, n=7623
Treg anchors, n=878
Tcon anchors, n=1325
p<2e-08
p<1e-43
LFC (Tcon ÷ Treg)
J
Loop Compartments
Proportion
Treg Anchors
Tcon Anchors
NS Anchors
p=1e-94
p=6e-134
More A in Tcon
More A in Treg
Difference in compartment score

**Figure S2: Quality, reproducibility and differential loop calling in Tcon and Treg Hi-C data. Related to Figure 1.**

(A) Number of contacts in our Tcon and Treg cell Hi-C data compared to data from Liu et al. 2023.

(B) Correlation of Hi-C contacts within Treg replicates (red), within Tcon replicates (blue) and between Treg/Tcon replicates (gray), at different distances from the diagonal.

(C) Histogram of Mustache FDR values for loops and inclusion criterion (FDR < .02).

(D) Fraction of TSSs, binned into five quantiles by RPKM, overlapping a loop anchor. Mann-Whitney U test (***, p < .001).

(E) MA plot for differential Hi-C analysis in loops using DESeq2.

(F) Venn diagram of loop anchors for loops that were Treg-specific, Tcon-specific or not significantly differential (NS) between Tcon and Treg, at a log2 fold change (LFC) threshold of 0.

(G) Pileups of Tcon-specific (left), non-differential (center), or Treg-specific (right) loops, in Treg Hi-C data (top) or Tcon Hi-C data (bottom).

(H) Scatter plot of log2 fold change (LFC) of gene expression (RNA-seq) and LFC of chromatin looping (Hi-C) aggregated over all loops for each gene. Blue, red; significant differential expression (FDR < .05). Pearson correlation calculated over all expressed genes.

(I) CDF plot of RNA-seq LFCs (Tcon / Treg) for genes overlapping NS, Treg-specific or Tcon-specific loop anchors. Mann-Whitney U test was performed between Treg- or Tcon-specific anchors and the NS anchors.

(J) CDF plot of difference in A/B compartment scores between Treg and Tcon bins overlapping Treg-specific, Tcon-specific, and non-differential loop anchors. Mann-Whitney U test was performed between Treg- or Tcon-specific anchors and the NS anchors.

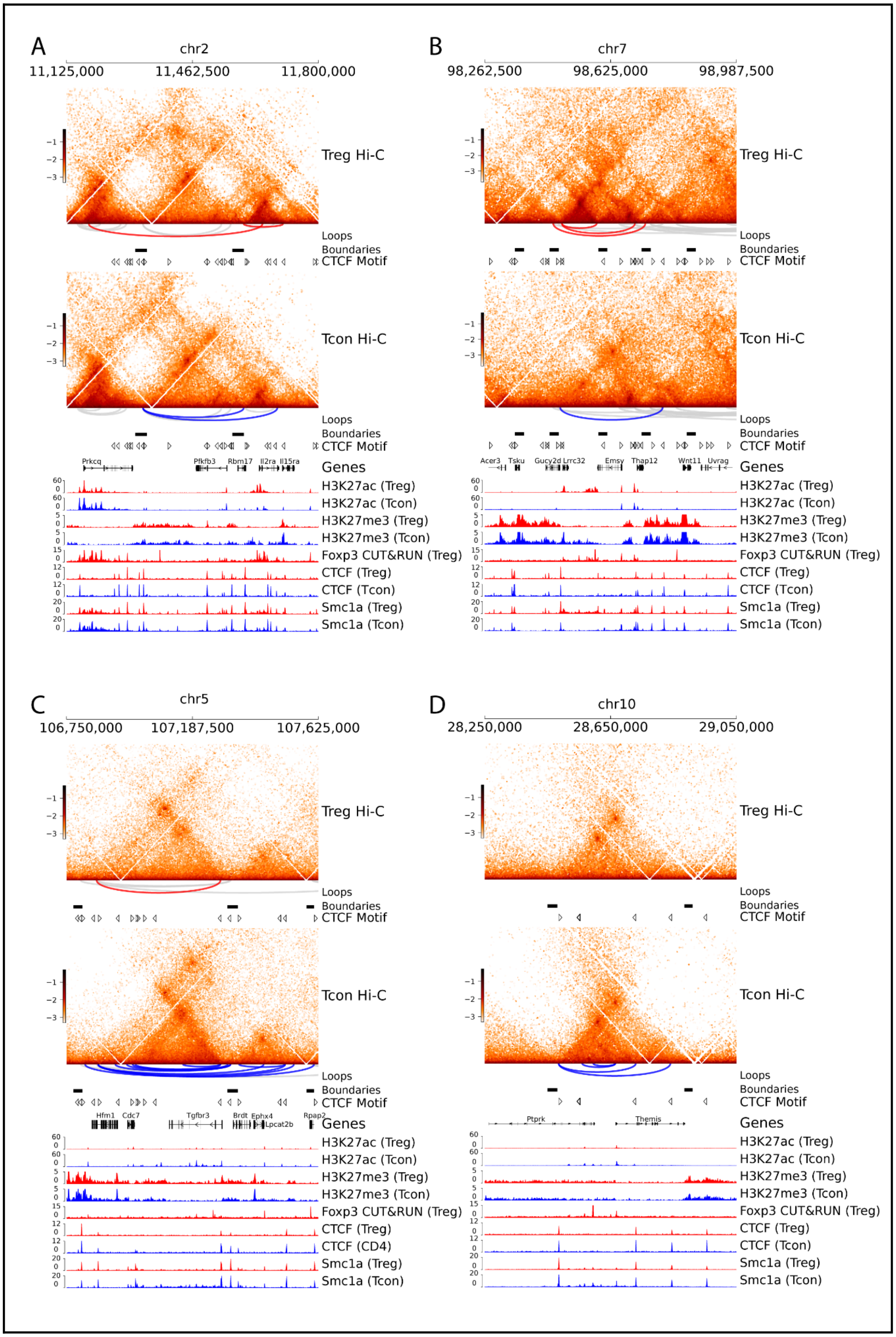
A
chr2
11,125,000
11,462,500
11,800,000
Treg Hi-C
Loops
Boundaries
CTCF Motif
Tcon Hi-C
Loops
Boundaries
CTCF Motif
Prkcq
Pfkfb3
Rbm17
Il2ra
Il15ra
Genes
H3K27ac (Treg)
H3K27ac (Tcon)
H3K27me3 (Treg)
H3K27me3 (Tcon)
Foxp3 CUT&RUN (Treg)
CTCF (Treg)
CTCF (Tcon)
Smc1a (Treg)
Smc1a (Tcon)
B
chr7
98,262,500
98,625,000
98,987,500
Treg Hi-C
Loops
Boundaries
CTCF Motif
Tcon Hi-C
Loops
Boundaries
CTCF Motif
Acer3
Tsku
Gucy2d
Lrrc32
Emsy
Thap12
Wnt11
Uvrag
Genes
H3K27ac (Treg)
H3K27ac (Tcon)
H3K27me3 (Treg)
H3K27me3 (Tcon)
Foxp3 CUT&RUN (Treg)
CTCF (Treg)
CTCF (Tcon)
Smc1a (Treg)
Smc1a (Tcon)
C
chr5
106,750,000
107,187,500
107,625,000
Treg Hi-C
Loops
Boundaries
CTCF Motif
Tcon Hi-C
Loops
Boundaries
CTCF Motif
Hfm1
Cdc7
Tgfbr3
Brdt
Ephx4
Lpcat2b
Rpap2
Genes
H3K27ac (Treg)
H3K27ac (Tcon)
H3K27me3 (Treg)
H3K27me3 (Tcon)
Foxp3 CUT&RUN (Treg)
CTCF (Treg)
CTCF (CD4)
Smc1a (Treg)
Smc1a (Tcon)
D
chr10
28,250,000
28,650,000
29,050,000
Treg Hi-C
Loops
Boundaries
CTCF Motif
Tcon Hi-C
Loops
Boundaries
CTCF Motif
Ptprk
Themis
Genes
H3K27ac (Treg)
H3K27ac (Tcon)
H3K27me3 (Treg)
H3K27me3 (Tcon)
Foxp3 CUT&RUN (Treg)
CTCF (Treg)
CTCF (Tcon)
Smc1a (Treg)
Smc1a (Tcon)

**Figure S3: Examples of Treg- and Tcon-specific loops and TADs. Related to Figure 1.**

Balanced Hi-C chromatin contact frequency and epigenomic tracks at the (A) *Il2ra* locus, (B) *Lrrc32* locus (C) *Tgfbr3* locus, and (D) *Themis* locus. Red arcs: Treg-specific loops, blue arcs: Tcon-specific loops, gray arcs: shared loops.

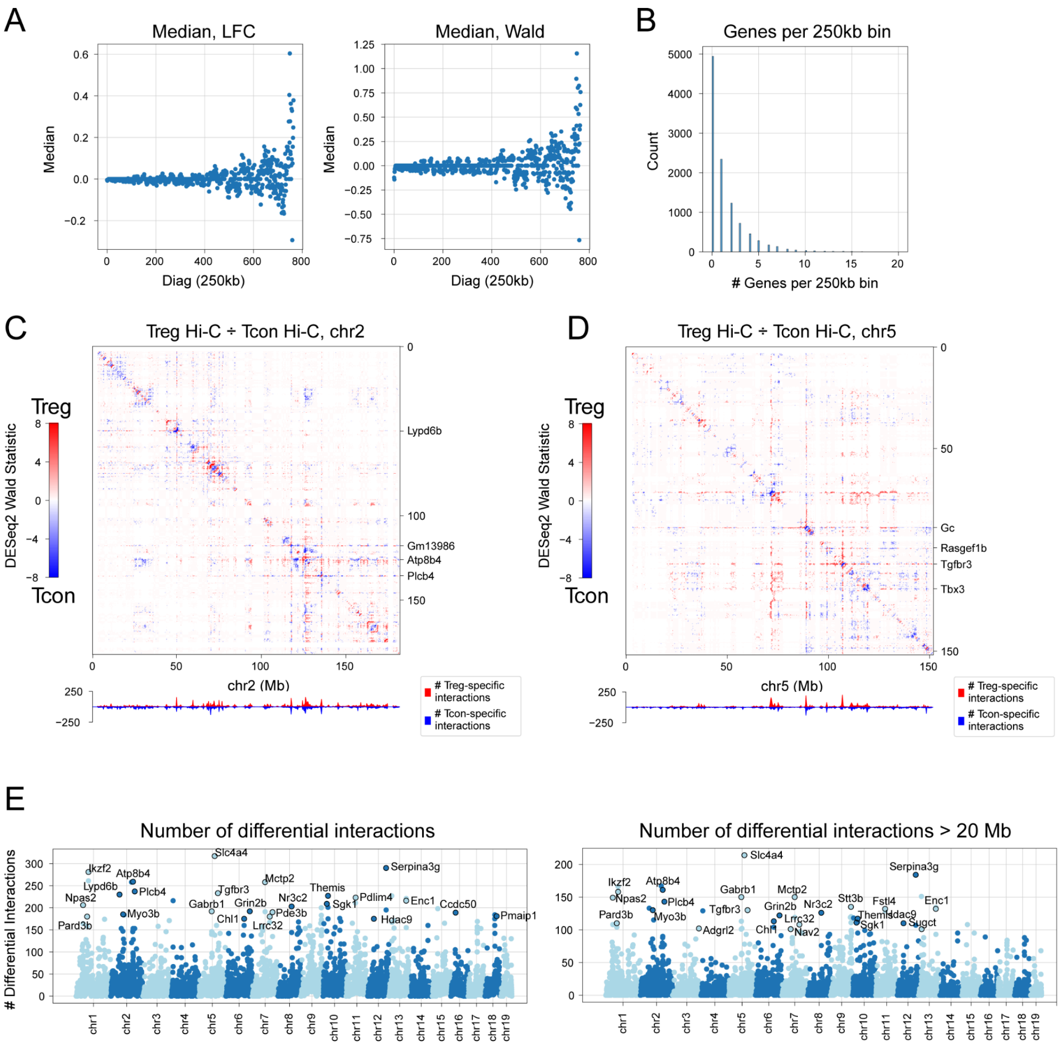


**Figure S4: Chromosome-wide differential Hi-C analysis at 250 Kb resolution. Related to Figure 1.**

(A) Scatter plot of median LFC (left) and DESeq2 Wald statistic (right) for all Hi-C interactions at a fixed genomic distance (x-axis), for genome-wide intrachromosomal differential Hi-C analysis at 250 Kb resolution between Tcon and Treg cells.

(B) Number of genes contained in each 250 Kb bin (excluding bins with no genes).

(C) Differential Hi-C analysis (using DESeq2) for 250 Kb genomic bins across chromosome 2. Analogous to Figure 1C.

(D) Differential Hi-C analysis (using DESeq2) for 250 Kb genomic bins across chromosome 5. Analogous to Figure 1C.

(E) Manhattan-style plot of the number of statistically significant differential Hi-C interactions between Tcon and Treg cells for each 250 Kb bin for all interactions (left) or for interactions at distances > 20 Mb (right).

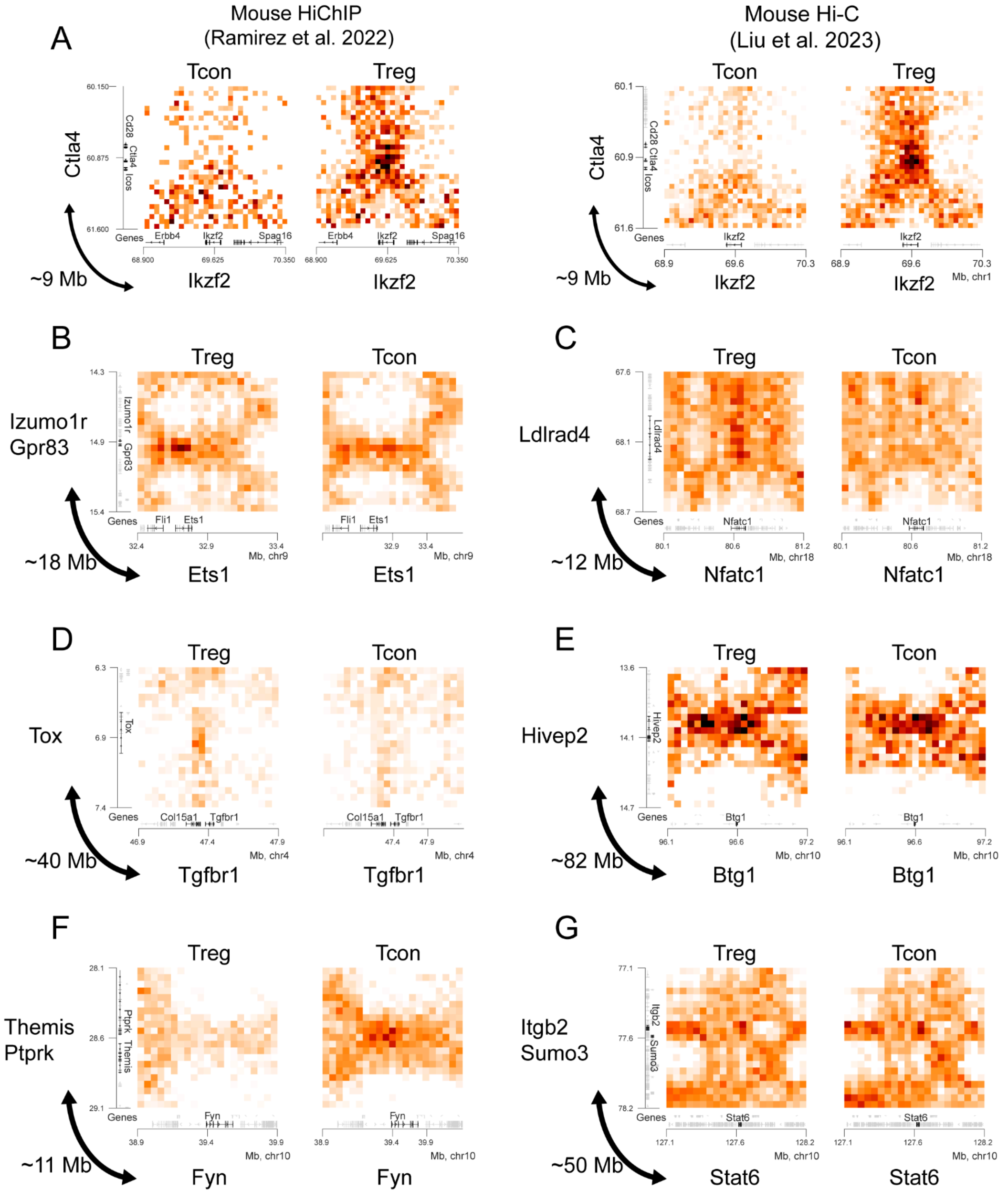


**Figure S5: Examples of distal metadomain genome interactions in Tcon and Treg cells. Related to Figure 3.**

(A) Balanced HiChIP contact maps at 50 Kb resolution for the interaction between Ikzf2 and the Cd28/Ctla4/Icos locus for data from Ramirez et al. 2022 (left) or data from Liu et al. 2023 (right). (B) Balanced Hi-C contact maps at 50 Kb resolution between Ets1 and Izumo1r, (C) Ldlrad4 and Nfatc1, (D) Tox and Tgfbr1, (E) Hivep2 and Btg1, (F) Themis and Fyn, and (G) Itgb2 and Stat6.

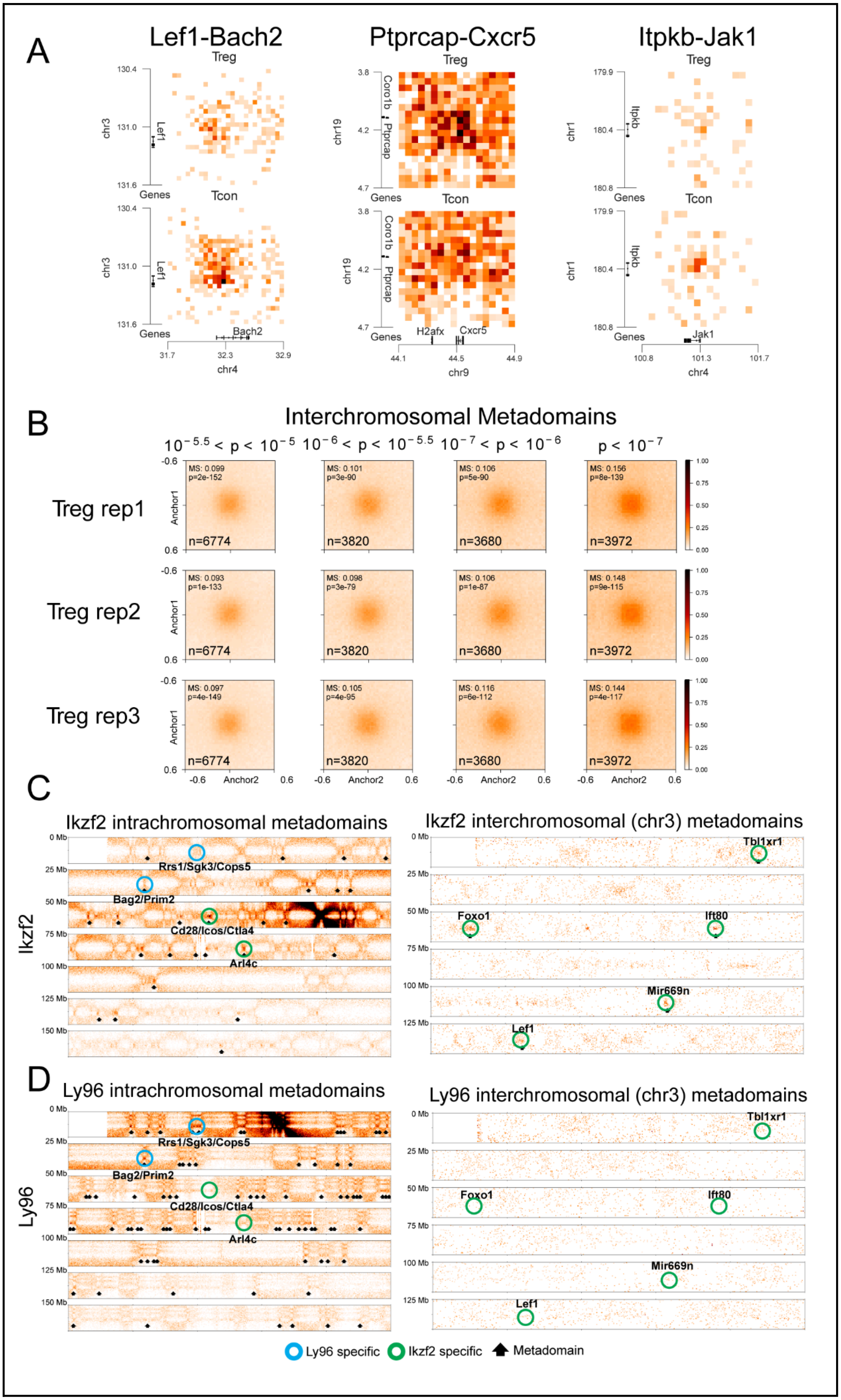
A
Lef1-Bach2
Ptprcap-Cxcr5
Itpkb-Jak1
Treg
Tcon
Genes
Lef1
Bach2
Coro1b
Ptprcap
H2afx
Cxcr5
Itpkb
Jak1
chr3
chr4
chr19
chr9
chr1
B
Interchromosomal Metadomains
$10^{-5.5} < p < 10^{-5}$
$10^{-6} < p < 10^{-5.5}$
$10^{-7} < p < 10^{-6}$
$p < 10^{-7}$
Treg rep1
Treg rep2
Treg rep3
Anchor1
Anchor2
n=6774
n=3820
n=3680
n=3972
C
Ikzf2 intrachromosomal metadomains
Ikzf2 interchromosomal (chr3) metadomains
Ikzf2
Rrs1/Sgk3/Cops5
Bag2/Prim2
Cd28/Icos/Ctla4
Arl4c
Tbl1xr1
Foxo1
Ift80
Mir669n
Lef1
D
Ly96 intrachromosomal metadomains
Ly96 interchromosomal (chr3) metadomains
Ly96
Ly96 specific
Ikzf2 specific
Metadomain

**Figure S6: Interchromosomal metadomains in Tcon and Treg cells. Related to Figure 3.**

(A) Balanced Hi-C data at 50 Kb resolutions for the metadomains linking Lef1 and Bach2, Ptprcap and Cxcr5, and Itpkb and Jak1. Multiple genes were omitted from genomic tracks for clarity.

(B) Pileup of interchromosomal Treg metadomains, by replicate, at different InterDomain p-value cutoffs.

(C) Hi-C signal for intrachromosomal (left) and interchromosomal (right; chromosome 3) interactions for the bin containing Ikzf2. Metadomains are annotated with a black arrow.

(D) Hi-C signal for intrachromosomal (left) and interchromosomal (right; chromosome 3) interactions for the bin containing Ly96. Metadomains are annotated with a black arrow.

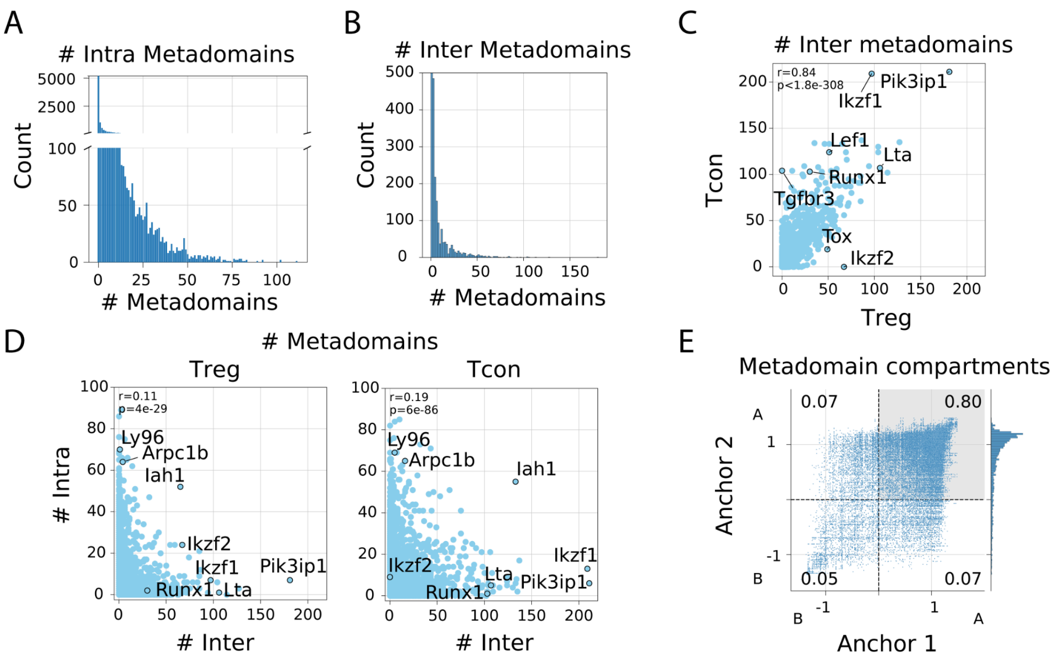


**Figure S7: Additional metadomain analysis. Related to Figure 3.**

(A) Histogram of the number of intrachromosomal metadomains called in Treg cells.

(B) Histogram of the number of interchromosomal metadomains called in Treg cells.

(C) Scatterplot of the number of interchromosomal metadomains per 250 Kb bin in Treg vs Tcon cells, for bins with more than 10 intra- or interchromosomal metadomains. Pearson correlation calculated over all bins.

(D) Scatterplot of the number of intrachromosomal metadomains per 250 Kb bin versus the number of interchromosomal metadomains per 250 Kb bin for all bins, in Treg cells (left) or Tcon cells (right). Pearson correlation calculated over all bins.

(E) Scatter plot of the A/B compartment score (at 250 Kb resolution) in Treg cells for genomic regions involved in metadomains.

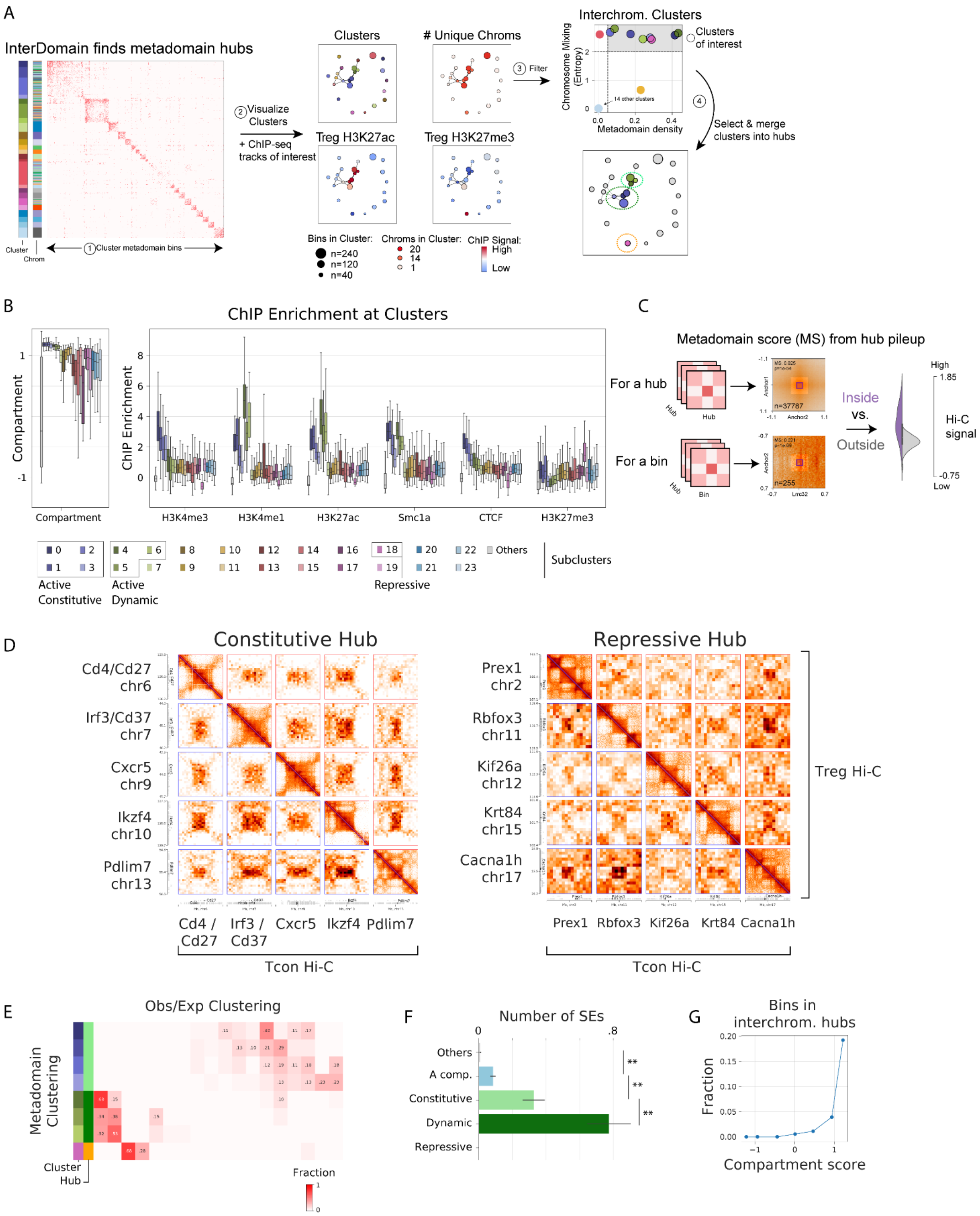


**Figure S8: Metadomain hubs. Related to Figure 4.**

(A) Identification of interchromosomal metadomain hubs. The 250 Kb bins with a large number of intrachromosomal or interchromosomal metadomains are clustered using hierarchical clustering. Interchromosomal clusters are selected using metadomain density and Shannon entropy of representation across chromosomes (see Methods). Clusters which share metadomains are then merged into interchromosomal hubs.

(B) Boxplots for normalized ChIP-seq signal and A/B compartment scores for 250 Kb bins in metadomain clusters.

(C) Schematic for the definition of the metadomain score (MS) representing the degree of interaction between all genomic bins within a metadomain hub, or between a single genomic bin and a metadomain hub. For this analysis, the corresponding interactions are selected (between all bin pairs in the hub, top; or between a single bin and the hub, bottom), and the balanced Log(O/E) Hi-C signal is calculated for each interaction (250 x 250 Kb centered at the interaction, “inside” values) and a flanking region (here, 900 x 900 Kb excluding the 250 x 250 Kb centered at the interaction, “outside” values). Distributions of these values (purple, gray) are then compared using the Mann-Whitney U test, and the metadomain score is defined as the difference of the means of these distributions. This analysis is visualized as a Hi-C pileup signal.

(D) Balanced Hi-C data (50 Kb resolution; upper triangle, Treg; lower triangle, Tcon) for selected genes from the Constitutive hub (left) and the Repressive hub (right).

(E) Robustness analysis for the definition of metadomain hubs. Clustering of the same 250 Kb bins as in panel B with the same procedure and number of clusters, but using all pairwise Obs/Exp Hi-C, instead of presence/absence of metadomains. Bins corresponding to the three interchromosomal metadomain hubs from Figure 4 were selected, and their cluster assignments in the Obs/Exp Hi-C clustering were visualized. Shown is the fraction of each new cluster overlapping with each metadomain cluster.

(F) Number of SEs overlapping bins in each hub. (**, $p < 1e-3$, Fisher’s Exact Test).

(G) Fraction of all bins in interchromosomal metadomain hubs, grouped by compartment score.

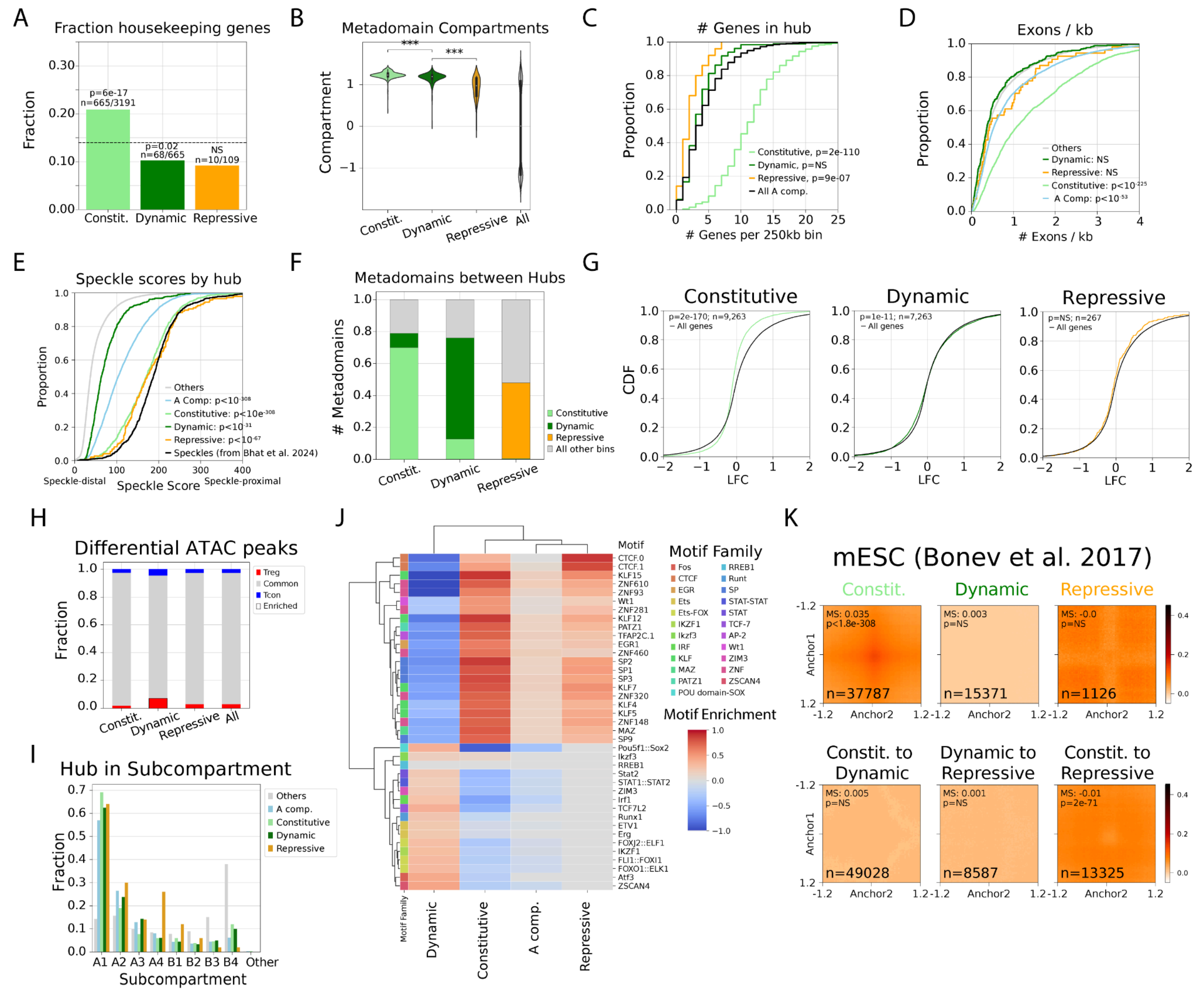


**Figure S9: Properties of metadomain hubs in Tcon and Treg cells. Related to Figure 4.**

(A) Fraction of housekeeping genes in each metadomain hub. Housekeeping gene annotations are from He and Williams 2020. Fisher's exact test.

(B) Violin plot for the distribution of the A/B compartment score in 250 Kb bins in metadomain hubs, compared to all other bins. Mann-Whitney U test (***, $p < .001$).

(C) Cumulative distribution function (CDF) plot for the number of genes per 250 Kb bin in metadomain hubs, compared to all bins in the A compartment. Mann-Whitney U test.

(D) CDF plot for the number of exons per Kb of transcript for genes in metadomain hubs, compared to genes in other bins. Mann-Whitney U test.

(E) CDF plot for speckle scores from Bhat et al. 2024 for bins in metadomain hubs, compared to other bins. Mann-Whitney U test.

(F) Barplot showing metadomain interactions from each metadomain hub to itself, to other metadomain hubs, and to bins outside of metadomain hubs.

(G) Differential accessibility analysis in metadomain hubs for ATAC-seq data for Tcon and Treg cells from van der Veeken et al. 2020. CDF plots for ATAC-seq LFCs (Treg / Tcon) for all peaks in each metadomain hub vs. all other peaks (Kolmogorov-Smirnov test).

(H) Quantification of the number of significantly differentially accessible (FDR < .05, |LFC| > .5) ATAC-seq peaks in each metadomain hub (The enrichment is significant in the Dynamic hub, Fisher's exact test).

(I) Fraction of each interchromosomal hub overlapping each subcompartment called in Yin et al. 2023.

(J) Motif enrichment for all ATAC-seq peaks in each metadomain hub. Motif enrichment is calculated as the LFC in frequency of the motif for peaks in the hub compared to the frequency of the motif in all other peaks. Non-significant motifs (Fisher's exact test, p > .05) are shown in grey. The ten most significant motifs were taken from each metadomain hub for visualization.

(K) Pileup of mESC Hi-C data from Bonev et al. 2017 for the three metadomain hubs defined in the Tcon/Treg cell Hi-C data. Tick marks are in Mb.

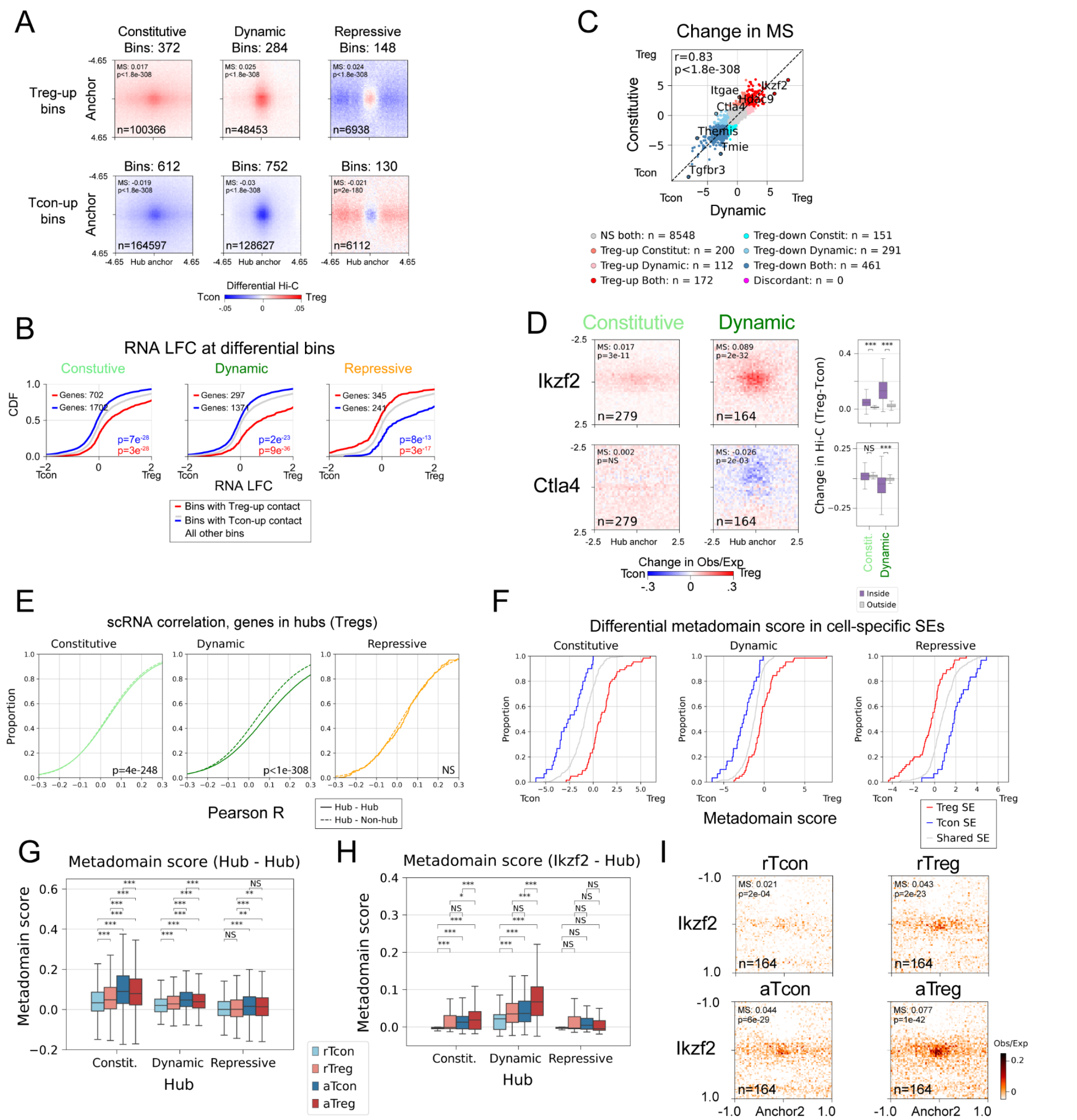


**Figure S10: Cell type specificity and differential gene expression in metadomain hubs. Related to Figure 5.**

(A) Pileup of Tcon and Treg Hi-C (25 Kb resolution) for 250 Kb genomic bins detected as involved in metadomain hubs in Treg- or Tcon-specific manner using differential metadomain score analysis.

(B) LFC (Treg vs. Tcon) of RNA-seq signal at 250 Kb bins with Treg- and Tcon-specific contact with the metadomain hubs. Mann-Whitney U test.

(C) A scatterplot of differential metadomain scores for interactions of 250 Kb genomic regions with the Active Constitutive and Active Dynamic metadomain hubs. Pearson correlation calculated over all bins.

(D) Differential Hi-C signal (25 Kb resolution) for interactions of *Ikzf2* and *Ctla4* with the active metadomain hubs. Boxplots (right) show quantification of the metadomain scores for these interactions. Boxplots: center line, median; box limits, upper and lower quartiles; whiskers, 1.5x interquartile range.

(E) CDF plots of Pearson correlation in Treg scRNA-seq gene expression data from Hemmers et al. 2019 for pairs of genes in the same metadomain hub (solid lines), or pairs of genes where one gene is in a metadomain hub and the other gene is not in the hub (dashed lines). Mann-Whitney U test.

(F) Differential metadomain scores between Treg and Tcon cells for Treg-specific SEs (red), Tcon-specific SEs (blue), or shared SEs (gray) taken from Kitagawa et al. 2017.

(G) Boxplot quantification showing metadomain scores for all bins in the metadomain hubs in rTcon, aTcon, rTreg and aTreg cells. Mann-Whitney U test (***, $p < .001$, **, $p < .01$).

(H) Boxplot quantification of metadomain scores between Ikzf2 and all other bins in the metadomain hubs in rTcon, aTcon, rTreg and aTreg cells. Mann-Whitney U test (***, $p < .001$, **, $p < .01$).

(I) Interchromosomal Hi-C pileups (25 Kb resolution) in active and resting Tcon and Treg cells showing interactions between Ikzf2 and the Active Dynamic hub.

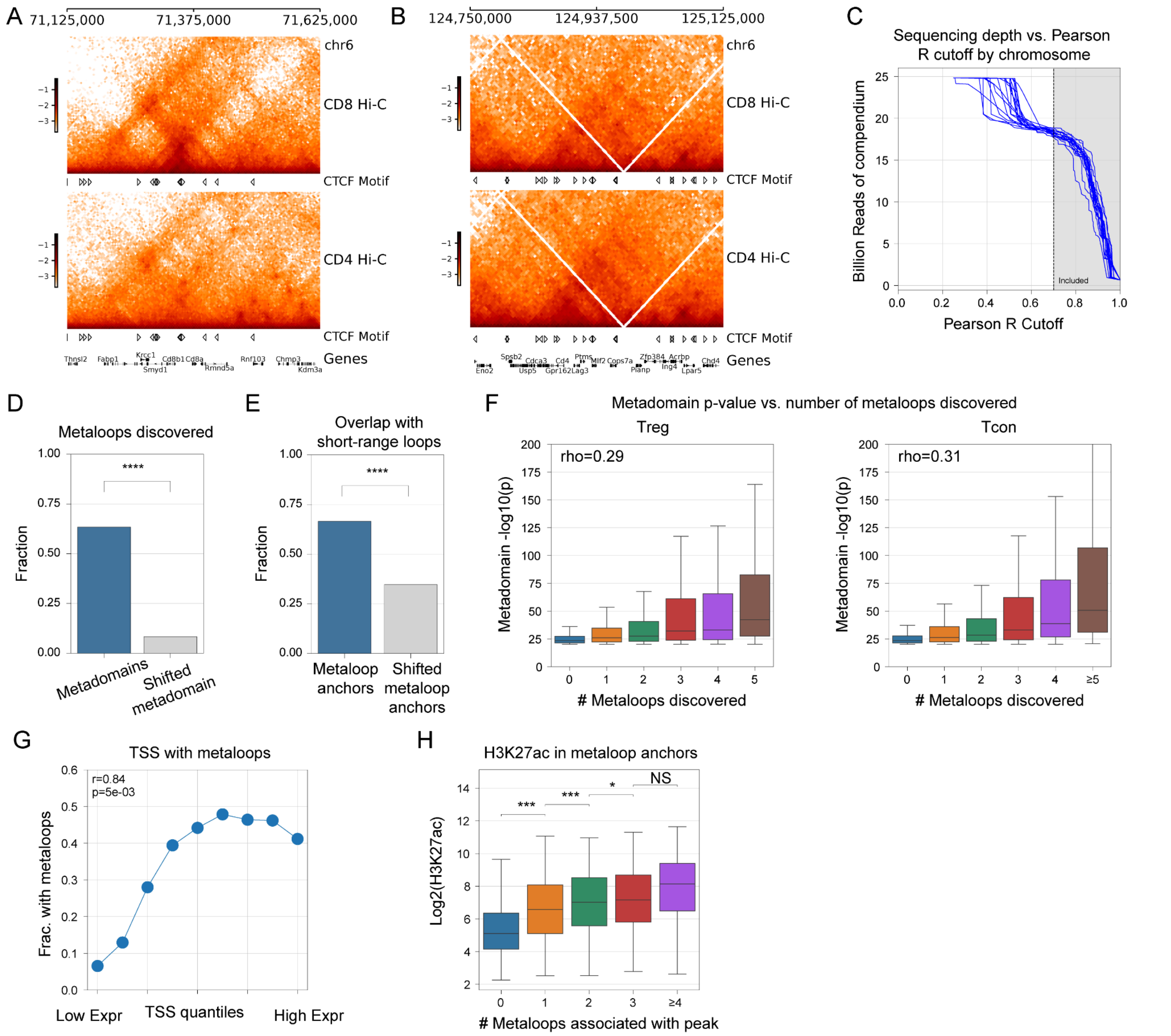


**Figure S11: T cell Hi-C compendium preprocessing and analysis of metaloop calls. Related to Figure 6.**

(A) Balanced CD8 and CD4 T cell Hi-C chromatin contact frequency and epigenomic tracks at the *Cd8a/Cd8b1* locus for the newly generated Hi-C data.

(B) Balanced CD8 and CD4 T cell Hi-C chromatin contact frequency and epigenomic tracks at the *Cd4* locus for the newly generated Hi-C data.

(C) Total sequencing depth as a function of Pearson R threshold used for including relevant Hi-C datasets.

(D) Fraction of metadomains in which metaloops could be detected at 5 Kb resolution in the T cell Hi-C compendium, compared with flanking regions (shifted by 250 Kb) (****, $p < 1e-5$, Fisher's Exact test).

(E) Fraction of anchors of metaloops called in the T cell Hi-C compendium that overlapped short-range loop anchors at 5 Kb resolution in Treg and Tcon data, compared with flanking regions (shifted by 50 Kb). (****, $p < 1e\text{-}5$, Fisher's Exact test).

(F) Boxplot of the Treg (left) or Tcon (right) -log10 InterDomain p-value for the original metadomain (y-axis) as a function of the number of metaloops recovered in that metadomain after metaloop identification in the T cell Hi-C compendium (x-axis). Spearman rho calculated over all points (not just boxplot medians).

(G) Fraction of TSSs engaging in metaloops, as a function of gene expression (RPKM).

(H) H3K27ac ChIP-seq signal as a function of the number of metaloops formed by each H3K27ac peak. (***, $p < 1e\text{-}3$; *, $p < .05$; Mann-Whitney U test)

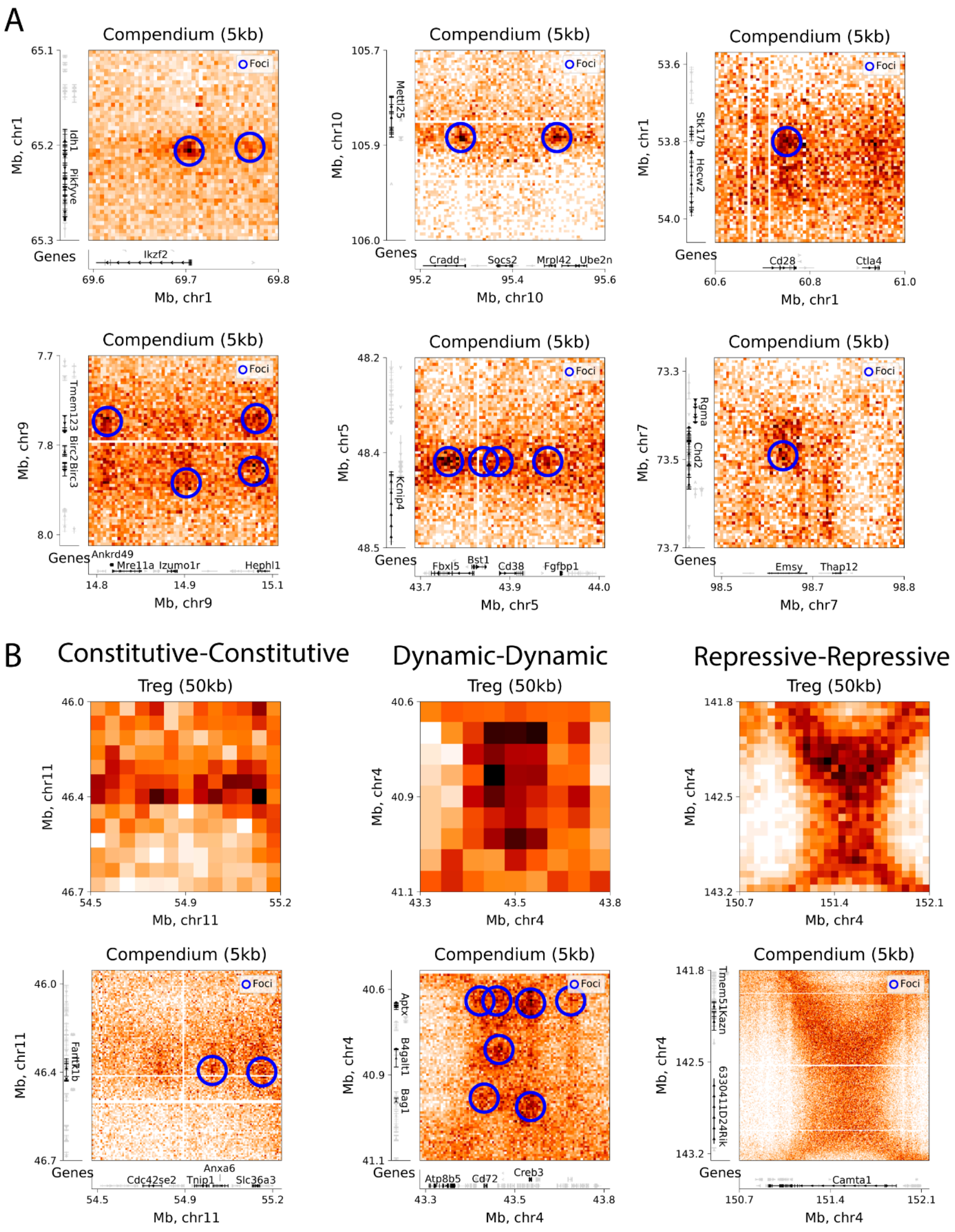


**Figure S12: Examples of metaloops in the T cell Hi-C compendium. Related to Figure 6.**

(A) Balanced Hi-C data at 5 Kb resolution in the T-cell Hi-C compendium for metadomains linking *Ikzf2* and *Idh1*, *Mettl25* and the *Socs2* locus, *Stk17b* and *Cd28*, *Izumo1r* and *Birc3*, *Cd38* and *Kcnjp4*, and *Emsy* and *Chd2*. Blue circles denote metaloops detected using InterDomain.

(B) (top) Balanced Hi-C data at 50kb resolution in Treg cells for metadomains linking bins in each metadomain hub. (bottom) Balanced Hi-C data at 5 Kb resolution in the T-cell Hi-C compendium for the same sites. Blue circles denote metaloops detected using InterDomain.

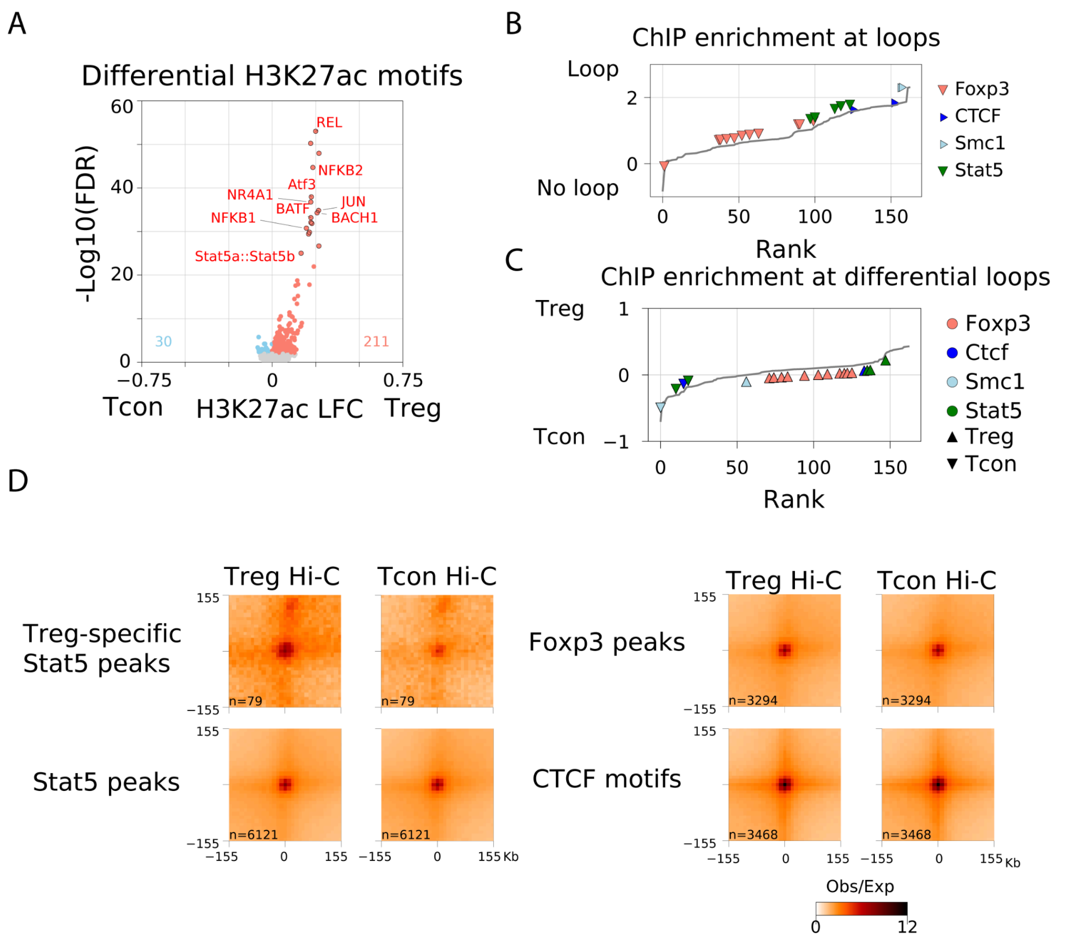


**Figure S13: T cell epigenomic compendium and TF motif analysis. Related to Figure 7.**

(A) Motif enrichment in Treg-specific vs. Tcon-specific H3K27ac peaks. LFC is calculated from the motif frequency in Treg-specific peaks / Tcon-specific anchors. FDR calculated using Fisher's exact test.

(B) Mean ChIP-seq signal in 5 Kb loop anchor loci vs. non-loop-anchor 5 Kb loci.

(C) Mean difference in ChIP-seq signal between Treg-specific loop anchors and Tcon-specific loop anchors. Upward arrows represent ChIP-seq data generated for Treg cells, while downward arrows represent ChIP-seq data for Tcon cells.

(D) Balanced O/E Hi-C pileup at 5 Kb resolution for loops whose anchors contain a Treg-specific Stat5 peak, Foxp3 peak, all Stat5 peaks, or a CTCF motif. Tick marks are in Kb.
